# Mapping spatial gradients in spatial transcriptomics data with score matching

**DOI:** 10.1101/2025.11.24.690257

**Authors:** An Wang, Donald Geman, Uthsav Chitra, Laurent Younes

## Abstract

Spatial transcriptomics (ST) technologies measure gene expression at thousands of locations within a two-dimensional tissue slice, enabling the study of spatial gene expression patterns. Spatial variation in gene expression is characterized by *spatial gradients*, or the collection of vector fields describing the direction and magnitude in which the expression of each gene increases. However, the few existing methods that learn spatial gradients from ST data either make restrictive and unrealistic assumptions on the structure of the spatial gradients or do not accurately model discrete transcript locations/counts. We introduce SLOPER (for Score-based Learning Of Poisson-modeled Expression Rates), a generative model for learning spatial gradients (vector fields) from ST data. SLOPER models the spatial distribution of mRNA transcripts with an *inhomogeneous Poisson point process (IPPP)* and uses *score matching* to learn spatial gradients for each gene. SLOPER utilizes the learned spatial gradients in a novel diffusion-based sampling approach to enhance the spatial coherence and specificity of the observed gene expression measurements. We demonstrate that the spatial gradients and enhanced gene expression representations learned by SLOPER leads to more accurate identification of tissue organization, spatially variable gene modules, and continuous axes of spatial variation (isodepth) compared to existing methods.

**Software availability:** SLOPER is available at https://github.com/chitra-lab/SLOPER.

## 1 Introduction

Spatial transcriptomics (ST) technologies make high-throughput measurements of gene expression at thousands of locations inside a two-dimensional (2-D) tissue slice. These technologies enable the study of spatial gene expression patterns across many biological systems including the brain, developing organs, and diseased tissues [1–5].

Broadly, the goal of spatial transcriptomics data analysis is to characterize spatial variation in gene expression, which reflects both global tissue organization and continuous variation in cell state. Tissues are organized into *spatial domains*, or discrete regions with distinct cell type composition and function (e.g. the different layers of the cerebral cortex), with many genes displaying large changes in expression near domain boundaries. Many genes also exhibit *continuous variation* in expression in specific directions due to local gradations in cell state or microenvironment; e.g., differentiation-related genes continuously vary along the crypt-villus axis of intestinal glands due to stem cell migration [6]. While numerous computational methods have been developed to identify spatial domains [7–16] and/or spatially variable genes and gene modules [12, 17–29], many of these methods do not explicitly model continuous variation in gene expression nor describe the direction in which gene expression varies.

The continuous variation of a function defined on a spatial domain is captured by its (spatial) gradient, which is the vector, evaluated at each point, formed by the function’s partial derivatives with respect to its input. In the context of ST, the *spatial gradient* for each gene is a *vector field* over the 2-D tissue slice where each vector indicates the direction and magnitude in which expression is increasing. Importantly, the collection of spatial gradients across *all* genes describes both spatial domains and continuous variation in gene expression: spatial domains are marked by genes with large spatial gradient vectors on the domain boundary, while continuous variation is indicated by smoothly varying spatial gradient vectors within or across domains. Thus, the task of learning spatial gradients from ST data is central to characterizing spatial patterns of gene expression.

Several recent computational methods aim to learn spatial gradients by computing the gradient of a function describing the (mean) expression at each spatial location. Direct estimation of such a gene expression function is challenging due to the low sequence coverage (large data sparsity) of current ST technologies, and so existing methods often place simplifying structural assumptions on this function. The common approach is to model gene expression as a function of a 1-D spatial coordinate that describes the relative position of a location in the tissue slice. While most approaches for estimating such a 1-D coordinate require substantial prior knowledge on the tissue geometry [30–33], one of the authors recently developed the GASTON suite of deep learning algorithms which learn a 1-D “isodepth” coordinate without such prior knowledge [34, 35]. However, a key assumption of the GASTON algorithms—and implicitly, all methods which model gene expression using a 1-D coordinate—is that the spatial gradient vectors for *every* gene point in the same direction, which is a highly restrictive and unrealistic assumption as different genes may exhibit spatial gradients in different directions; e.g. in the intestine, genes vary along both the crypt-villus and proximal-distal axes [6, 36]. One exception is the recently introduced StarTrail algorithm [37] which does not use a 1-D coordinate parametrization and instead models gene expression using a Gaussian process which is highly inefficient to fit to data. Moreover, Gaussian processes and other spatial autocorrelation approaches [28] do not directly model the *discrete* ST measurements (i.e., mRNA transcript locations or counts), instead *smoothing* the sparse and spatially diffuse gene expression and potentially masking underlying spatial patterns [38]. Thus, to the best of our knowledge, there are no methods that efficiently and accurately learn spatial gradients, nor use these gradients to characterize spatial expression patterns.

*Score matching methods*, introduced in [39] and recently adapted to generative artificial intelligence (AI) models [40, 41], learn the gradient of a log-probability density function, also called the *score*. Their key advantage is that estimating this gradient bypasses the need to compute normalizing constants of probability distributions and allows for the estimation of models that can be much more complex than those accessible through traditional likelihood-based methods. Moreover, the estimated gradient is often combined with diffusion-based sampling methods such as Langevin dynamics [40, 41], allowing for the generation of random samples from the underlying distribution. The broad success of score matching in generative modeling of complex and diverse data (e.g. images, videos) suggests a potential application towards learning and utilizing spatial gradients in ST data.

In this paper, we introduce SLOPER (for Score-based Learning Of Poisson-modeled Expression Rates), a score-based algorithm for accurately and efficiently estimating spatial gradients of gene expression from ST data. SLOPER models spatial gene expression using an *inhomogeneous Poisson point process (IPPP)* model which describes the spatial distribution of mRNA transcripts across a tissue, and uses score matching to efficiently learn the spatial gradient (i.e. the *score*) for each individual gene. SLOPER leverages the estimated spatial gradients in a novel *annealed Langevin dynamics* formulation to generate *“enhanced”* gene expression representations that are sharper and more spatially coherent than the original gene expression measurements. We demonstrate that the spatial gradients and enhanced gene expression learned by SLOPER enable many downstream analyses including the identification of tissue organization (spatial domains) and distinct spatial patterns of gene expression (gene modules). Furthermore, we derive an algorithm to learn a 1-D isodepth coordinate [34, 35] from the SLOPER-estimated spatial gradients, demonstrating SLOPER’s improved generalizability compared to existing 1-D coordinate ST methods. Using 10x Genomics Visium ST data from the human dorsolateral prefrontal cortex (DLPFC) and Slide-SeqV2 ST data from the mouse testis, we demonstrate that the SLOPER more accurately learns spatial domains, gene modules, and isodepth compared to existing methods. In particular, SLOPER reveals spatial colocalization patterns of testicular cell types at different temporal stages of spermatogenesis.

## 2 Methods

### 2.1 An inhomogeneous Poisson point process model of spatial gene expression

We derive a generative model of the spatial distribution of the expression of each gene measured using spatial transcriptomics (ST) technologies. For a fixed gene, ST technologies measure the spatial location **s**_1_ = (*x*_1_, *y*_1_), …, **s**_*N*_ = (*x*_*N*_, *y*_*N*_) of *N* mRNA transcripts in a 2-D tissue slice *T* ⊆ ℝ^2^. We assume that transcript locations **s** = (*x, y*) may assume arbitrary locations across the tissue slice *T*, which is true for ST technologies with sub-cellular spatial resolution such as MERFISH [42] or Xenium [43]. In Section 2.2, we discuss how to extend our approach to ST technologies with lower spatial resolution (e.g. 10x Genomics Visium [44], Slide-Seq [45, 46]) where one measures aggregated transcript counts in segmented cells or on a fixed grid rather than individual transcript locations.

We model the distribution of the *N* measured transcript locations 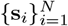 for a given gene with an *inhomogeneous Poisson point process (IPPP)*. Poisson point processes are a classical and well-studied statistical model for the distribution of points in space, where the *inhomogeneity* allows for non-constant transcript densities across space [47–49]. The IPPP is parametrized by an *intensity function λ* : *T* → [0, ∞), where *λ*(*x, y*) quantifies the local expected density of transcripts at spatial position (*x, y*) ∈ *T*. For a given region *R* ⊆ *T* (representing, e.g., a single cell), the number *N*(*R*) = |{*i* : *s*_*i*_ ∈ *R*}| of observed transcripts in is a random variable following a Poisson distribution: *N* (*R*) ∼ Poisson (Λ(*R*)), where Λ(*R*) = ∫ _(*x,y*) ∈ *R*_ *λ*(*x, y*) *dx dy* = *E*[*N* (*R*)] is the average number of transcripts in the region *R*. Under the IPPP, conditioned on the total number *N* = *N* (*T*) of transcripts for the given gene in the tissue slice *T*, then the individual transcript locations **s**_1_, …, **s**_*N*_ are independently and identically distributed (IID) within *T* with probability density function (PDF) 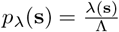, where we define Λ = Λ(*T*).

The intensity function *λ* reflects both technical and biological variation in the distribution of mRNA transcripts. For example, the number of mRNA transcripts measured in a cell may vary for technical reasons including sequencing depth, capture efficiency and misannotated cell boundaries. Many existing single-cell and spatial analyses account for such technical variation through *“normalization”*, where one divides measured gene expression counts by a factor such as the total number of mRNA inside a cell (*“library size normalization”*) or the volume of a cell [50–52]. In a similar manner, we incorporate both technical and biological variation into our model by assuming that the intensity function *λ* factorizes as:

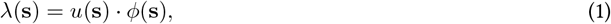

where the term *u*(·) : ℝ^2^ → ℝ models spatial variation in *technical* factors, such as the total number of transcripts or cell volume, and *ϕ*(**s**) : ℝ^2^ → ℝ is the *normalized* intensity function which describes biological variation in mRNA transcript distribution. When it is clear from context, we may omit the term “normalized” and directly call *ϕ* the intensity function. In our analyses, we set the technical factor *u*(**s**) equal to the total number of measured mRNAs inside the cell *C* that contains spatial location **s** = (*x, y*), which intuitively corresponds to the library size normalization used in single-cell RNA sequencing analysis, although one may also set the technical factor term *u*(**s**) equal to other quantities such as the total volume of a cell [52].

A *spatial gradient* describes spatial variation in gene expression across the tissue slice *T*. Here, we define the spatial gradient as the gradient ∇ log *ϕ* of the *log* of the (normalized) intensity function *ϕ*. Note that the gradient ∇ log *ϕ* of the log-intensity is *scale-invariant*, i.e., ∇ log *ϕ* = ∇ log(*cϕ*) for any positive scalar *c >* 0.

Given observed transcript locations 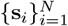, we aim to learn the spatial gradient ∇ log *ϕ*. We formalize this as the following computational problem.

#### Spatial Gradient Problem (SGP)

Let **s**_1_ = (*x*_1_, *y*_1_), …, **s**_*N*_ = (*x*_*N*_, *y*_*N*_) be spatial locations sampled from an inhomogeneous Poisson point process in a tissue slice *T* with intensity function *λ*(**s**) = *u*(**s**) · *ϕ*(**s**). Given spatial locations 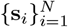 and technical factor term *u*(**s**), compute the spatial gradient ∇ log *ϕ*.

### 2.2 A score-based method to estimate spatial gradients

We propose to solve the SGP using *score matching* [39]. Score matching is a popular approach for directly estimating the *score* ∇_*s*_ log *p*(**s**) of a distribution *p*(**s**), in contrast to likelihood-based methods which aim to directly estimate the distribution *p*(**s**) [39–41]. We use score matching because the distribution 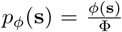, defined over spatial locations **s ∈** *T* in the tissue slice, has score equal to the spatial gradient we wish to compute in the SGP, i.e., ∇_*s*_ log *p*_*ϕ*_(*s*) = ∇_*s*_ log *ϕ*(*s*). We call *p*_*ϕ*_ the *normalized* distribution as it corresponds to the normalized intensity function *ϕ*, in contrast to the *observed* distribution *p*_*λ*_(**s**) which is the distribution of observed spatial locations **s**_1_, …, **s**_*N*_ drawn from the IPPP (Section 2.1).

#### Score matching and truncated score matching

In score matching, one aims to learn a *score function f*_*θ*_ : *T* → ℝ^2^ (typically a neural network) with parameters *θ* that approximate the spatial gradient ∇ log *ϕ*, i.e., *f*_*θ*_(**s**) ≈ ∇ log *ϕ*(**s**) at all spatial locations **s** ∈ *T*. This is accomplished by minimizing the score matching objective ℒ_SM_(*θ*) defined as:

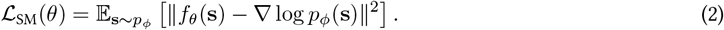

A major difficulty in minimizing ℒ_SM_ is that it involves the unknown distribution *p*_*ϕ*_. The key insight of the classical score matching methodology (e.g., the seminal paper by Hyvärinen [39]) is that under certain so-called “regularity” conditions, the score matching objective ℒ_SM_(*θ*) is equal to the following expression:

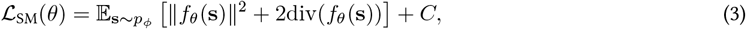

where *C* is a constant that does not depend on *θ*.

In our ST setting, however, we cannot directly apply the classical score matching methodology. This is because *p*_*ϕ*_ is a *truncated* distribution defined on a bounded region *T* (i.e. the tissue slice), and recent work by Liu et al. [53] and Cao et al. [54] has shown that the regularity conditions used to derive (3) do not hold for truncated distributions. Instead, Liu et al. [53] propose minimizing the following *truncated* score matching objective

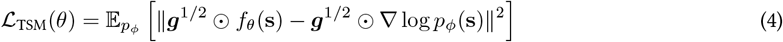

where ***g*** = [*g*_0_, *g*_0_] is a spatial weighting vector derived from the boundary distance function *g*_0_(**s**) = min_**z**∈*∂T*_ dist(**s, z**), where *∂T* is the boundary of the tissue domain *T* and ⊙ denotes the element-wise product. The boundary distance term ***g*** ensures that the objective ℒ_TSM_ satisfies the score matching regularity conditions. Liu et al. [53] show that, if *f*_*θ*_ is twice continuously differentiable and ∇ log *p*_*ϕ*_ is continuously differentiable, then the truncated score matching objective ℒ_TSM_(*θ*) is equal to

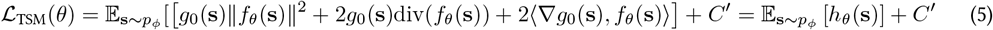

where we define *h*_*θ*_(**s**) = *g*_0_(**s**) ∥ *f*_*θ*_(**s**) ∥^2^ + 2*g*_0_(**s**)div(*f*_*θ*_(**s**)) + 2 ⟨∇*g*_0_(**s**), *f*_*θ*_(**s**) ⟩, and *C*^*′*^ is a constant that does not depend on *θ*.

#### Importance reweighting

In score matching, the expectation 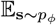 is replaced by empirical averages over training data. However, in our setting, observations do not follow *p*_*ϕ*_, but the observed distribution *p*_*λ*_. We address this issue using *importance reweighting* [55]. Specifically, we rewrite the expectation 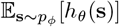 as an expectation over the observed distribution *p*_*λ*_:

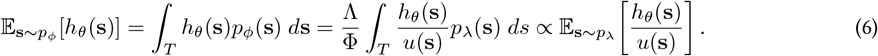

Since the proportionality coefficient does not depend on *θ*, (5) can be minimized using the r.h.s. of this equation. In practice, since we are given the observed spatial locations **s**_1_, …, **s**_*N*_ and the technical factor term *u*(**s**) (Section 2.1), we can replace 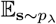 an empirical average and solve the following optimization problem:

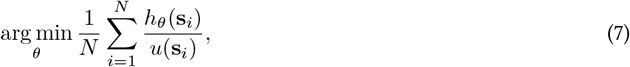

#### 2.2.1 Technologies with aggregated spatial measurements

The proposed optimization problem in (7) assumes that we know the location **s**_*i*_ of each individual transcript. However, many spatial transcriptomics technologies (e.g. 10x Genomics Visium and Visium HD [44] and Slide-SeqV2 [45, 46]) only report *aggregated* measurements over spatially localized regions, such as spots or segmented cells. These ST technologies partition the tissue *T* = *C*_1_ ∪ *C*_2_ ∪ · · · *C*_*K*_ into *K* disjoint spatial regions *C*_*k*_ ⊆ ℝ^2^, where each region *C*_*k*_ corresponds to an individual cell or spot, and report the centroid coordinate **c**_*k*_ ℝ^2^ and the number *a*_*k*_ = |{*i* : **s**i ∈ *C*_*k*_}| of transcripts located in each region *C*_*k*_. The individual transcript locations **s**_*i*_ are unknown.

We propose a simple approach for adapting the score matching optimization problem (7) to aggregated transcript count measurements. Under the mild assumption that each region *C*_*k*_ is sufficiently small and the objective function *h*_*θ*_ (**s**) varies smoothly within the region *C*_*k*_, then average value 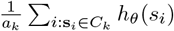 of the objective function over transcripts in region *C*_*k*_ is approximately equal to its value *h*_*θ*_(**c**_*k*_) at the centroid **c**_*k*_:

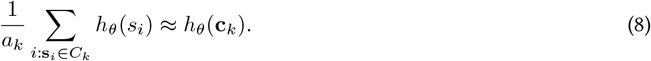

Substituting (8) into the truncated score matching objective (7) and simplifying yields the following problem:

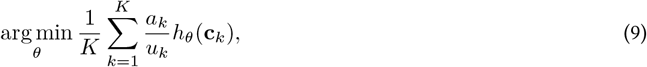

where *u*_*k*_ is the total number of mRNA molecules measured in region *k*.

### 2.3 Enhancing gene expression with annealed Langevin dynamics

A key benefit of score matching methods is that the score function can be used in Markov chain Monte-Carlo (MCMC) simulations to generate random samples that approximately match the observed data. Given the score ∇ log *ϕ* of a distribution *p*_*ϕ*_, the Langevin dynamics, provided by the stochastic differential equation (SDE)

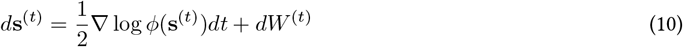

is (under some hypotheses) such that **s**_*t*_ approximately follows *p*_*ϕ*_ for large *t*, where *W* ^(*t*)^ is a Brownian motion [56]. For example, AI-based generative models use a variant of this principle for approximately sampling from *p*_*ϕ*_ [40, 41]. However, we empirically observe that directly sampling from the distribution *p*_*ϕ*_(**s**) often produces diffuse and noisy spatial patterns—due to the stochasticity and low counts of the observed ST data—that are difficult to utilize in downstream analyses.

Instead, we propose an approach for producing *enhanced* gene expression measurements using the learned scores ∇ log *ϕ*, by simply sampling from a distribution proportional to 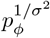. When *σ* is small, sampling from 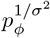 produces more samples from the high-probability regions in the tissue slice *T* (i.e. large *p*_*ϕ*_) and less samples from low-probability regions. Our sampling approach is obtained by modifying (10) to

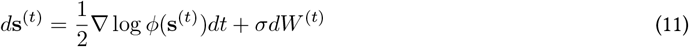

and the limit distribution provides a crisper, more interpretable, view of the intensity function *ϕ* associated with each gene. To accelerate the convergence of this algorithm, we will use an annealed dynamic in which we let *σ* depend on time; that is, we take taking *σ*_*t*_ = *α*^*t*^ for some *α <* 1. Numerically, this SDE can be discretized using an Euler-Maruyama scheme [57], leading to iterations

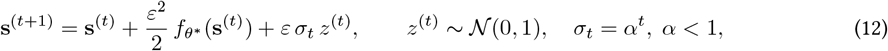

for discrete times *t*, where *ε >* 0 controls the drift step size and *H* is the number of time steps.

We run these iterations for a large number *H* of steps (i.e. *t* = 1, …, *H*) under *N* random initial conditions 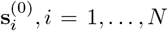, *N* to generate a set 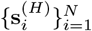 of transcript locations for the gene expression, which we call the *Langevin-enhanced* transcript locations. See Supplement for details on parameter selection.

Since many ST technologies partition the tissue into pre-defined regions 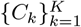 corresponding to individual cells or spots (Section 2.2.1), we additionally *bin* the enhanced coordinates 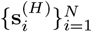 into the regions 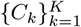. Specifically, we compute the *Langevin-enhanced* gene expression vector 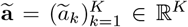 where entry 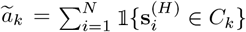 is the enhanced transcript count for region *C*_*k*_. This binning step improves the interpretability of our enhanced gene expression locations and enables their use in downstream analysis.

### 2.4 SLOPER algorithm and applications

We implement the truncated score matching (Section 2.2) and annealed Langevin dynamics (Section 2.3) methods in a package called SLOPER (Fig. 1), available at https://github.com/chitra-lab/SLOPER.

**Figure 1:**
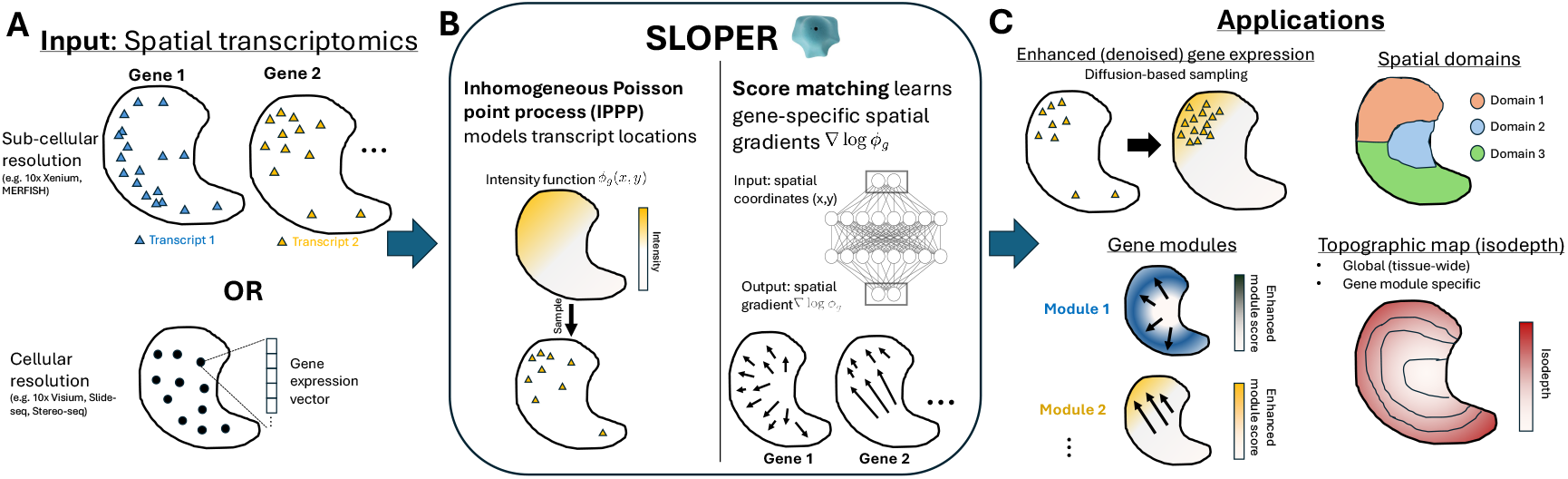
SLOPER learns spatial gradients of gene expression with score matching. **(A)** The input to SLOPER is ST data from a single 2-D tissue slice, either at sub-cellular resolution (transcript locations) or cellular resolution (gene expression vectors). **(B)** SLOPER models transcript locations / gene expression counts with an inhomogeneous Poisson point process (IPPP) model, which is parametrized by a (normalized) intensity function *ϕ*_*g*_ (*x, y*) for each gene *g*. SLOPER uses a deep neural network to directly estimate the spatial gradient (or *score*) ∇log *ϕ*_*g*_ for each gene. **(C)** The spatial gradients learned by SLOPER have several applications including enhancing the spatial coherence and specificity of gene expression using diffusion-based sampling (annealed Langevin dynamics); the identification of spatial domains and (spatially variable) gene modules; and the inference of global and module-specific topographic maps defined by 1-D isodepth coordinates.

Given ST data consisting of either transcript locations or gene expression vectors for *G* genes (Fig. 1A), we model the transcript distribution for each gene *g* = 1, …, *G* according to an IPPP distribution with (normalized) intensity function *ϕ*_*g*_. SLOPER first minimizes the truncated score matching objective (7), parametrizing the score function 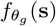 for each gene *g* with a neural network of two residual blocks [58], with Fourier feature positional encoding of the spatial coordinates **s** = (*x, y*) following [59]. We optimize the parameters *θ*_*g*_ of the neural network using the Adam optimizer [60], yielding an estimate 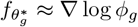 of the score (spatial gradient) for each gene *g* = 1, …, *G* (Fig. 1B). Subsequently, given a partition *T* = *C*_1_ ∪ · · · ∪ *C*_*K*_ of the tissue slice (e.g. segmented cells or spots; Section 2.2.1), SLOPER uses the estimated spatial gradient to generate a *Langevin-enhanced* gene expression vector **a**_*g*_ ∈ ℝ^*K*^for each gene *g* = 1, …, *G* (Section 2.3). See Supplement for details on hyperparameter selection.

The spatial gradients ∇log *ϕ*_*g*_ estimated by SLOPER are broadly useful for characterizing tissue organization and spatial variation in gene expression. We demonstrate their utility by showcasing several downstream tasks enabled by the SLOPER-estimated spatial gradients (Fig. 1C) including the enhanced gene expression **a** and the identification of spatial domains, spatially variable gene modules, and 1-D spatial axes of variation.

#### 2.4.1 Spatial domain identification

A key task in spatial transcriptomics data analysis is the identification of *spatial domains*, or regions *R* ⊆ *T* of the tissue slice *T* containing cells/spots with similar gene expression profiles and whose boundaries have large changes in gene expression (i.e. large spatial gradients). While it may be possible to infer spatial domains directly from the SLOPER-estimated spatial gradients 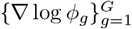, we instead propose a conceptually simple and efficient approach for domain identification using the Langevin-enhanced gene expression vectors **a**_*g*_ (Section 2.3), whose entries concentrate around high-probability regions of the density *p*_*ϕ*_—that is, spatial domains. Specifically, we cluster the *Langevin-enhanced* transcript count matrix **A** = [**a**_*g*_] ∈ ℝ^*K×G*^, whose columns are the enhanced gene expression vector **a**_*g*_ for each gene *g* = 1, …, *G*, by first applying standard single-cell data normalization to the enhanced count matrix **A** (per-cell UMI normalization and log-transformation as implemented in Scanpy [61]) and then performing *k*-means clustering.

#### 2.4.2 Gene modules

We identify *gene modules*, or groups of (spatially variable) genes with similar expression patterns, by clustering the gene-specific spatial gradients ∇ log *ϕ*_*g*_ learned by SLOPER. Specifically, we define a similarity metric *ρ*(*g, g*^*′*^) between a pair *g, g*^*′*^ of genes as the average *cosine similarity* between their respective spatial gradients ∇ log *ϕ*_*g*_(**s**), ∇ log *ϕ*_*g*_*′* (**s**) across the tissue slice *T* :

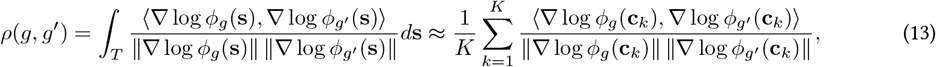

where 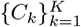 is a partition of the tissue and **c**_*k*_ is the centroid of region *C*_*k*_ (Section 2.2.1). We perform spectral clustering [62] using the similarity metric *ρ* to estimate *L* gene modules, where the number *L* of modules is specified in advance. We emphasize that in contrast to popular approaches for identifying gene modules (e.g. [28]) which define the similarity *ρ* between two genes using the observed gene expression—which may be spatially diffuse and noisy—we define the gene-gene similarity *ρ* using their *spatial gradients*, which score matching accurately learns.

We further construct an interpretable, per-cell *gene module score* for each inferred module ℳ ⊆ {1, …, *G*} which summarizes the spatial expression pattern of the genes g ∈ ℳ in the module. To construct our score, we first define the module-ℳ normalized intensity function *ϕ*_ℳ_ ≜ Π_*g* ∈*ℳ*_ *ϕ*_*g*_(**s**)^1*/*|ℳ|^ as the geometric mean of the normalized intensity functions {*ϕ*_*g*_}_*g* ∈ ℳ_ for all genes *g* ∈ ℳ; equivalently, the module-ℳ spatial gradient 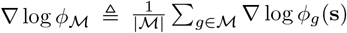 is equal to the arithmetic mean of the gene-specific spatial gradients {∇ log *ϕ*_*g*_}_*g*∈*M*_. Given spatial gradients {∇ log *ϕ*_*g*_}_*g*∈ℳ_ estimated by SLOPER, we run the annealed Langevin dynamics (Section 2.3) using the module-ℳ spatial gradient ∇ log *ϕ*_ℳ_; the module score is given by the resulting transcript locations/counts.

#### 2.4.3 Isodepth (1-D spatial axis) inference

Several recent computational methods developed by one of the authors learn spatial gradients under the strict assumption that the spatial gradient for each gene is proportional to the gradient ∇*d*(*x, y*) of a single 1-D scalar function *d* : ℝ^2^ → ℝ which they call the *isodepth* [30, 34, 35]. Here, we show that the the SLOPER-estimated gene-specific spatial gradients ∇ log *ϕ*_*g*_ allow us to generalize this approach and estimate gene set-specific isodepths *d*_ℳ_ : ℝ^2^ → ℝ for *any* subset ℳ ⊆ {1, …, *G*} of genes (e.g. a gene module).

Briefly, we assume that the spatial gradient ∇ log *ϕ*_*g*_ for each gene *g* ∈ ℳ is proportional to the gradient ∇*d*_ℳ_ of a 1-D function *d*_ℳ_ : ℝ^2^ → ℝ. The key observation is that *gradient outer product (GOP) matrix* 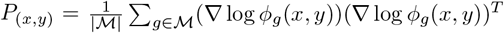, which may be computed from the SLOPER-estimated spatial gradients ∇ log *ϕ*_*g*_, provides a method for computing any *directional derivative* of the isodepth *d*_ℳ_. Specifically, we show that for any vector **v** ∈ ℝ^2^, the quantity 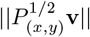 is proportional to the directional derivative of the isodepth *d*_ℳ_ at location (*x, y*) in the direction 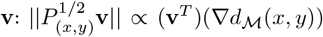. We then use the forward Euler method to estimate the function *d*_ℳ_ (*x, y*) given a single value *d*_ℳ_ (*x*_0_, *y*_0_) at a fixed point (*x*_0_, *y*_0_) and the directional derivative (∇*d*_ℳ_)^*T*^ **v** in any direction **v** ∈ ℝ^2^. See Supplement for more details.

## 3 Results

### 3.1 Simulations

We evaluated SLOPER on simulated IPPP data. Briefly, the intensity function *ϕ* is a weighted sum of three Gaussian ridge functions over a rectangular tissue *T* = [−1.5, 1.5] × [−1.5, 1.5], and is parametrized by a sparsity coefficient *c* ≥ 1 that controls the number of transcripts (smaller *c* yields sparser data). We divided the tissue *T* uniformly into *K* = 10, 000 spots 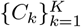 and sampled a transcript count *a*_*k*_ in each spot *k*. See Supplement for details.

We evaluate the performance of SLOPER’s score matching approach, which directly estimates the spatial gradient 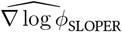, versus approaches which first estimate the intensity function *ϕ* and then compute its (log-) gradient. To this end, we compared SLOPER with a kernelized intensity estimator (KIE), a widely-used non-parametric approach to directly learn the intensity function of an IPPP [63]. We adapted the KIE to derive an estimate 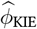 of the normalized intensity *ϕ* (see Supplement) and then computed the analytical gradient of 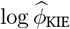. We measured accuracy using the *average cosine similarity* between the true spatial gradients ∇ log *ϕ* and the SLOPER- and KIE-estimated spatial gradients 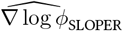 and 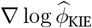, respectively; we only evaluate the cosine similarity on spatial locations (*x, y*) where the estimated KIE intensity 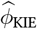 is non-zero to avoid division-by-zero errors. To account for the stochasticity of neural network training, we report the mean cosine similarity of SLOPER trained with five different random seeds.

We find that SLOPER more accurately estimates the spatial gradient ∇ log *ϕ* compared to the KIE across most parameter settings (Fig. 2A; Fig. S1). In particular, SLOPER consistently outperforms the KIE for less sparse data (i.e., larger sparsity coefficient *c*). Visually, we observe that compared to the ground truth spatial gradient (Fig. 2B), the SLOPER-estimated spatial gradient vector field 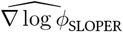 is more spatially coherent than the KIE-estimated gradient 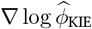 (Fig. 2C-D). Our results demonstrate that by directly learning a spatial gradient from data, SLOPER more accurately estimates the spatial gradient compared to traditional approaches which analytically compute a gradient of an estimated function.

**Figure 2:**
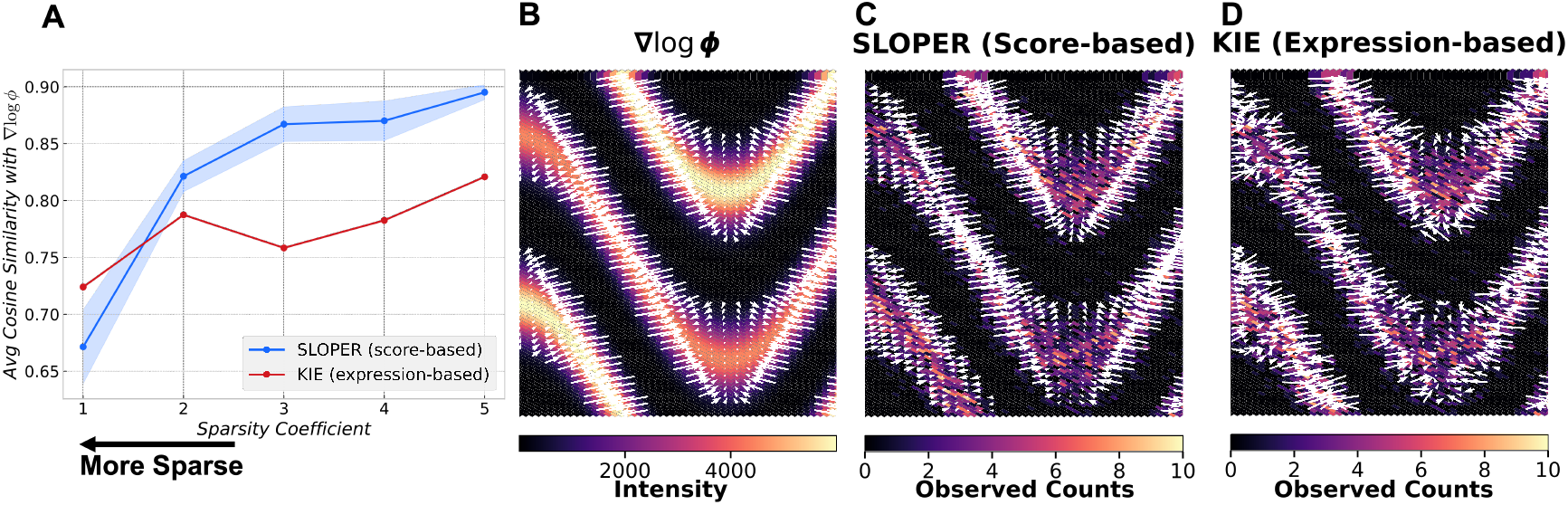
Benchmark of SLOPER and kernel intensity estimator (KIE) on simulated data. Only the left half of the tissue slice *T* is shown for clarity; results for the entire tissue slice *T* appear in Fig. S1B–D. **(A)** Average cosine similarity between the estimated and ground-truth spatial gradient ∇ log *ϕ* across sparsity coefficients *c* = 1, …, 5. Shaded regions denote the standard deviation across five random seeds for SLOPER. **(B)** Intensity function *ϕ* and spatial gradient ∇ log *ϕ*, shown with white arrows. **(C-D)** (C) SLOPER-estimated spatial gradient 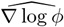 and (D) KIE-estimated spatial gradient 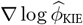, shown with white arrows, overlaid on the simulated Poisson counts *a*_*k*_.

### 3.2 Human dorsolateral prefrontal cortex (DLPFC)

We next evaluated SLOPER on a 10x Genomics Visium ST sample of the human dorsolateral prefrontal cortex (DLPFC) [64]. This dataset measures the expression of 33538 genes in 3639 cells. We used geneCover [65] to perform label-free marker selection and analyzed *G* = 500 genes. Each cell is manually annotated with one of the six cortical layers and white matter (WM) domains, providing a useful benchmark for spatial domain identification methods. We used SLOPER to learn spatial gradients and subsequently identify spatial domains, and compared the SLOPER-identified domains to spatial domains identified by (1) GraphST, a state-of-the-art domain identification method [15]; (2) domains identified by clustering the KIE-estimated intensity function 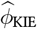; and (3) domains identified by standard single-cell clustering approaches, Leiden clustering [66] and *k*-means [67]. We had each method identify 7 domains to match the number of domains in the manual annotation, and we evaluated each method by computing the normalized mutual information (NMI) between the estimated domains and manually annotated layers. See Supplement for details.

SLOPER more accurately identifies spatial domains compared to GraphST, KIE, Leiden clustering, and *k*-means (Fig. 3A). Notably, SLOPER uniquely identifies Layer 2 (blue domain in Fig. 3B, SLOPER) compared to the other methods. In contrast, KIE, Leiden, and *k*-means fail to fully resolve Layers 3-5 (Fig. S2). The improved performance of SLOPER demonstrates the power of SLOPER’s score matching framework in modeling tissue organization.

**Figure 3:**
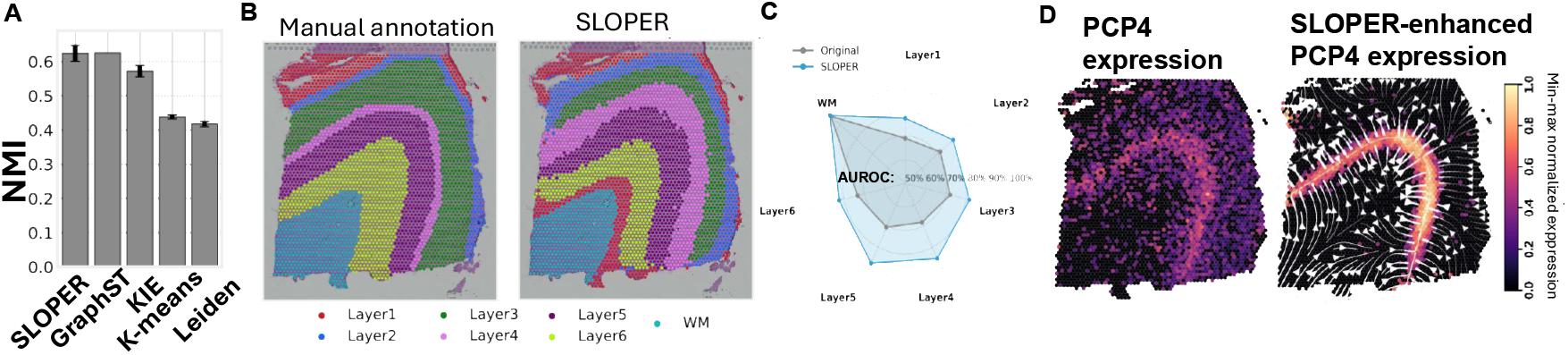
Benchmarking SLOPER domain detection and SLOPER-enhanced gene expression on DLPFC. **(A)** Normalized mutual information (NMI) of spatial domains identified by SLOPER and four other methods. **(B)** Left: manually annotated spatial domains [64]. Right: spatial domains identified by SLOPER. SLOPER domains are colored according to the manually annotated domain with largest overlap. **(C)** Comparison of average AUROC scores between original and SLOPER-enhanced gene expressions in identifying domain-specific genes from Layers 1–6 and white matter spatial domains. **(D)** Left: Min-max normalized *PCP4* expression. Right: Min-max normalized SLOPER-enhanced *PCP4* expression, where white streamlines show the learned gradient 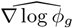.

A key distinguishing feature of SLOPER is its *Langevin-enhanced* gene expression, which reveals spatial patterns that are challenging to ascertain from the sparse and spatially diffuse observed gene expression values. As an illustration, we evaluated the performance of the original expression values **a**_*g*_ and the SLOPER Langevin-enhanced expression values 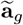 in determining the domain(s) of domain-specific genes *g*, quantified using an area under the receiver operating characteristic curve (AUROC) metric (see Supplement for details). The SLOPER Langevin-enhanced expression 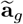 consistently achieves higher average AUROC than the original expression **a**_*g*_ across DLPFC Layers 1-6, with similar AUROC in the WM domain (Fig. 3C), indicating that SLOPER’s Langevin enhancement procedure improves the spatial coherence and specificity of domain-specific gene expression profiles. For example, *PCP4* is a well-established marker of DLPFC Layer 5 [64]. While *PCP4* exhibits relatively large expression values in other DLPFC domains (Fig. 3D left, S3A), the SLOPER-enhanced *PCP4* expression is sharply localized in DLPFC Layer 5 (Fig. S3B), accurately delineating the corresponding layer boundaries (Fig. 3D right). We observe similar enhancement of spatial specificity for other layer-specific and layer-enriched marker genes, including *KRT17, HPCAL1, VAMP1*, and *HOPX* (Fig. S4). These analyses demonstrate how SLOPER’s annealed Langevin dynamics “denoise” the observed gene expression profiles while preserving biological relevance.

We further demonstrate the versatility of SLOPER by using the SLOPER-estimated spatial gradients 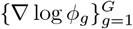 and a forward Euler method approach (Section 2.4.3) to compute the isodepth, a 1-D axis of spatial variation proposed by GASTON [34]. We compare the reference GASTON isodepth, which measures cortical layer depth in the DLPFC [34], to the SLOPER-estimated isodepth and the GraphST embeddings. We find (Fig. 4A) that SLOPER learns a 1-D isodepth coordinate (Fig. 4B) that closely matches the GASTON isodepth (Fig. 4C; Spearman *ρ* = 0.92). In contrast, the GraphST embeddings are not highly correlated with the isodepth (mean Spearman *ρ* = 0.09 across GraphST embedding components), with the closest matching GraphST embedding visually appearing spatially incoherent (Fig. 4D; Spearman correlation of 0.53 with the GASTON isodepth). These results demonstrate that the SLOPER-estimated spatial gradients provide an accurate, interpretable, and continuous representation of cortical layer organization. Beyond global structure, we further applied the isodepth inference to individual SLOPER gene modules (Fig. S5). The resulting module-specific isodepths reveal both layer-associated depth gradients (e.g., the isodepth of module 14 aligns with Layer 5) and non-laminar spatial topologies, such as the isodepth of module 1, which reflects the organization of spatially variable genes *SCGB2A2* and *SCGB1D2* [68].

**Figure 4:**
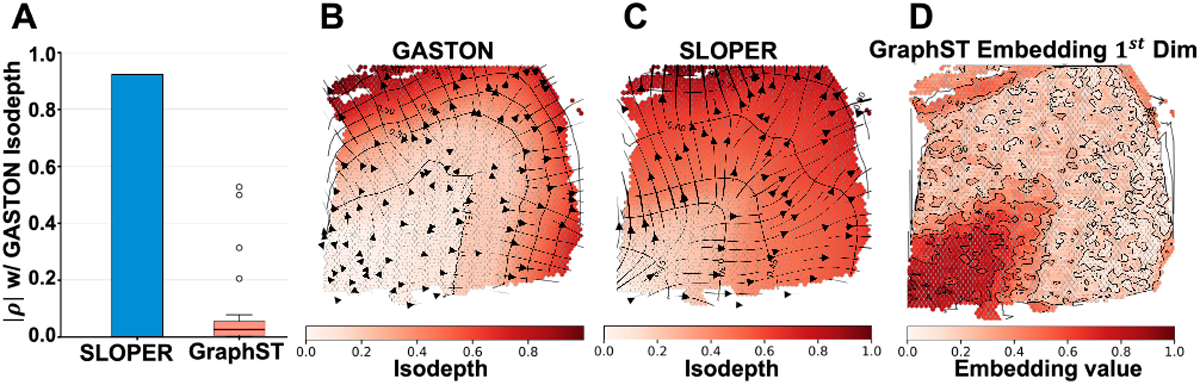
Benchmarking isodepth inference in the DLPFC. **(A)** Spearman’s correlation of inferred isodepths with the GASTON reference for SLOPER-inferred isodepth and GraphST embedding components. **(B–D)** Visualization of (B) GASTON and (C) SLOPER-inferred isodepth, and (D) GraphST embedding component with the highest Spearman correlation with the GASTON isodepth.

### 3.3 Mouse testis

We next applied SLOPER to the Slide-seq ST sample of the mouse testis [69]. This dataset measures the expression of 23705 genes in 39206 cells; as in the previous section, we performed label-free marker selection using geneCover [65] and analyzed 500 selected genes. The testis is composed of numerous repeating tubular structures in which spermatogenesis takes place, and we hypothesized that SLOPER would reveal subtle spatial gene expression patterns that reflect the spatial organization of the testis.

We compared the spatial domains identified by SLOPER to domains identified by GraphST, KIE, GASTON, and IRIS [13]. IRIS is a domain identification method which requires a single-cell reference dataset with labeled cell types and was previously applied to this mouse testis dataset. We had each method identify 5 domains, matching the number of IRIS domains in the original manuscript [13]. We visually observe (Fig. 5A) that the SLOPER-identified spatial domains that are substantially more similar to the IRIS-identified domains compared to the GraphST-, KIE-, and GASTON-inferred domains—despite the fact that SLOPER does not use an external single-cell reference while IRIS does. While there are no ground truth domain annotations for this dataset, IRIS performs extensive biological validation of their inferred domains, providing a biological interpretation for the different SLOPER-identified domains. For example, SLOPER and IRIS domain 1 (Fig. 5A) correspond to spermatocytes, a granular population colocalizing at the boundary of spermatids (SLOPER domains 2, 4 and IRIS domain 4, Fig. 5A). Importantly, GraphST, KIE, and GASTON do not identify the spermatocyte domain, with GASTON unable to identify any repeating domains (Fig. 5A). See Supplement and Fig. S6 for further analysis and interpretation of SLOPER domains.

**Figure 5:**
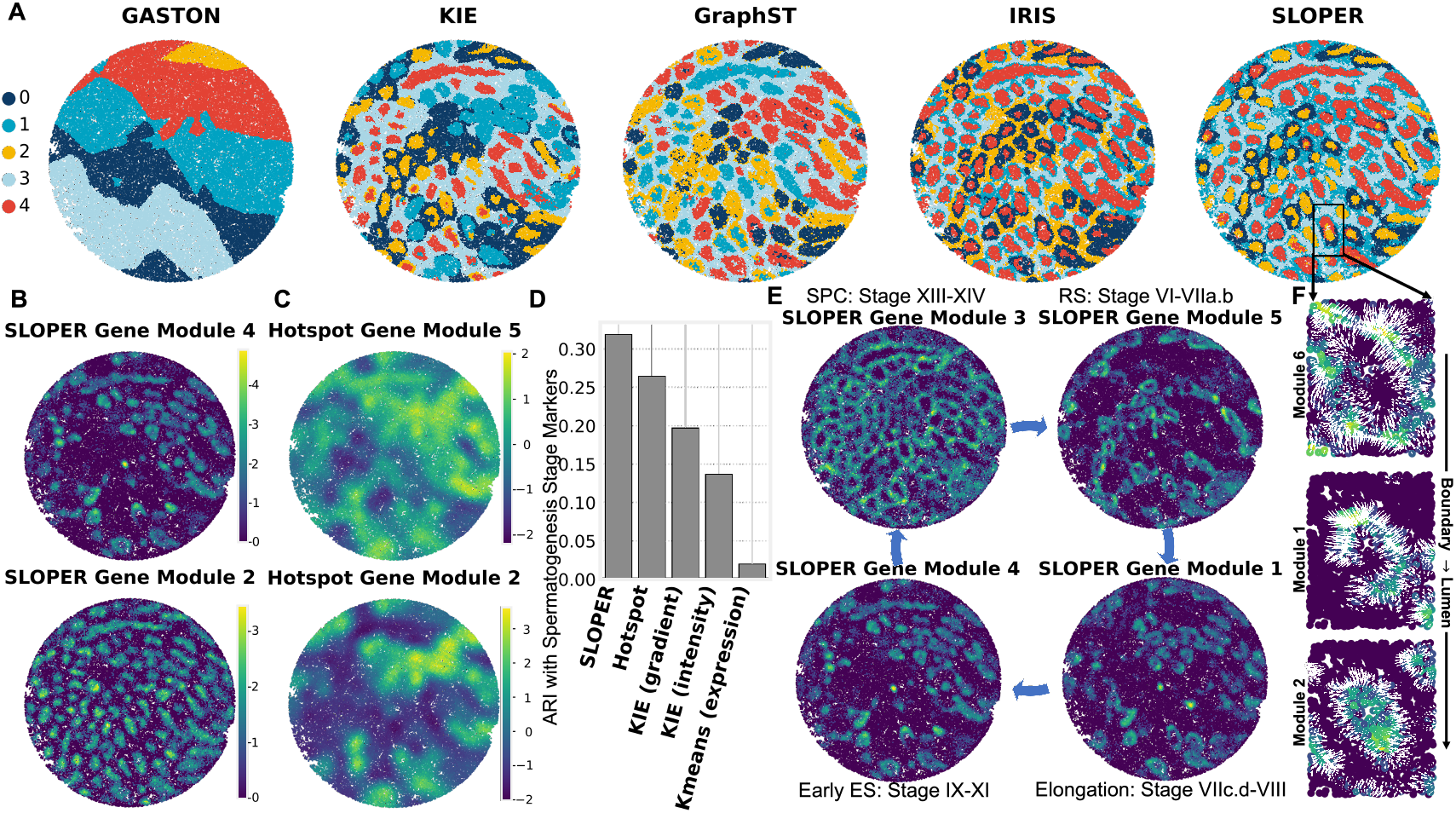
SLOPER identifies spatial domains and gene modules that demonstrate the spatiotemporal progression of cell types in the mouse testis. **(A)** Spatial domains identified by GASTON, IRIS, GraphST, KIE, and SLOPER. **(B)** SLOPER gene modules 4 (top) and 2 (bottom) colored by SLOPER module score. **(C)** Hotspot modules with the largest correlation with corresponding SLOPER module scores in (B), colored by Hotspot module score. **(D)** Adjusted Rand index (ARI) between known spermatogenesis stage marker genes [71] and gene modules identified by SLOPER and other approaches. **(E)** Selected SLOPER gene modules and spermatogenesis stages with highest marker gene overlap. Colors indicate SLOPER gene module score. Gene modules are ordered according to corresponding spermatogenesis stage. **(F)** SLOPER modules 6, 1, and 2, colored by SLOPER module score. Modules are ordered according to inferred spatiotemporal progression of spermatid maturation within the seminiferous tubule. White arrows indicate learned module-specific spatial gradient ∇ log *ϕ*_ℳ_ (Section 2.4.2).

A key distinction between SLOPER and IRIS is that SLOPER splits up IRIS domain 4—which represents elongated spermatids [13]—into two domains (SLOPER domains 2 and 4, Fig. 5A). We performed a differential expression analysis to assess whether these two SLOPER-identified elongated spermatid domains (Fig. S7A) reflect biologically distinct tubular structures in the testis. We computed the Wilcoxon rank-sum statistics for all genes and found extensive differential expression between the two domains (Fig. S7B). Interestingly, we observe a hierarchical pattern of differential expression, where genes differentially expressed in SLOPER domain 4 exhibit substantially lower expression in SLOPER domain 2 (e.g. *Prm1, Tnp1, Tnp2*; Fig. S8), while genes differentially expressed in SLOPER domain 2 exhibit fairly similar expression values in SLOPER domain 4 (e.g. *Prm2, Smcp, Pgk2*; Fig. S9). Indeed, prior biological studies [69, 70] found that the expression of SLOPER domain 4-specific genes *Prm1, Tnp1*, and *Tnp2* peak at earlier developmental stages than the expression of SLOPER domain 2-specific gene *Prm2*. These observations suggest that SLOPER domains 2 and 4 represent successive stages in the maturation of elongated spermatids, with domain 4 corresponding to early elongation and domain 2 to later-stage elongated spermatids. Importantly, SLOPER domains 2 and 4 were not identified by IRIS (Fig. 5A, IRIS) despite its use of an auxiliary reference dataset. Our results demonstrate the unique advantages of SLOPER’s Langevin dynamics framework in delineating subtle differences between spatial domains.

We next evaluated the gene modules and module scores computed by SLOPER. We compared SLOPER to Hotspot [28], a popular method for gene module identification; since Hotspot reports 10 gene modules, we used SLOPER’s spectral clustering approach to identify 10 modules. We observe that the SLOPER modules, represented by their module score (Methods, Section 2.4.2), resolve major testicular cell types more clearly than the Hotspot modules. For example, SLOPER module 4 and SLOPER module 2 visually correspond to SLOPER *domain* 4 and SLOPER domains 2, 4, respectively (Fig. 5B). (In the previous paragraph, we hypothesized that SLOPER domains 2 and 4 represent two populations of elongated spermatids at different developmental stages.) On the other hand, the corresponding Hotspot gene modules (i.e. Hotspot modules whose module scores are most correlated with the corresponding SLOPER module scores) appear spatially diffuse and oversmoothed, lacking the spatial resolution necessary to visually delineate the testicular spatial organization (Fig. 5C, S11). Moreover, as a validation of our domain-based differential expression analysis, we observe that SLOPER modules 2 and 4 contain *Prm1* and *Prm2* (Fig. S10A-B), respectively—genes that were differentially expressed in corresponding SLOPER domains 2 and 4, respectively (Fig. S8-S9). See Supplement for interpretation of other SLOPER modules.

We quantitatively validated the SLOPER gene modules using a list of known spermatogenesis developmental stage-specific marker genes [71]. Our validation is motivated by our observation that SLOPER module scores for modules 2 and 4 (Fig. 5B) concentrate on elongated spermatids at different developmental stages. We compared the SLOPER gene modules to those identified by Hotspot and three other baselines: (1) spectral clustering on the KIE-inferred gradient 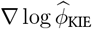, (2) *k*-means on genes represented by KIE-estimated normalized intensity 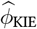, and (3) *k*-means on original expression profiles.

We find (Fig. 5D) that the SLOPER modules have a substantially larger ARI with the known spermatogenesis markers (ARI = 0.32) than the other approaches (ARI *<* 0.26). Furthermore, the overlap between SLOPER gene modules and the known developmental stage markers (Fig. S13A) and the SLOPER gene module scores reveal the spatial gene expression patterns for different developmental stages (Figure 5E). To illustrate, SLOPER modules 3, 9, and 10 (Fig. S12) show high overlap with spermatogenesis Stage XIII–XIV marker genes; e.g. SLOPER module 10 contains the canonical spermatocyte (SPC) marker gene *Aurka* [72]. Spermatocytes in stages XIII–XIV are enriched for pachytene spermatocytes and mark the completion of meiosis (Stages XII–XIV), giving rise to haploid round spermatids [71, 73]. Furthermore, the module score for SLOPER modules 3, 9, 10 is large in regions that visually correspond to the labeled spermatocyte cell types reported in the original Slide-seq study [69], suggesting that the spermatocytes cells identified in the original study are likely pachytene spermatocytes specifically. Through similar reasoning, we also find that SLOPER module 5 (Figure 5E, top right) consists of marker genes for round spermatids (RS) with elevated expression at Stage VI; SLOPER module 1 (Figure 5E, bottom right) consists of genes involved in the transition from round spermatids toward elongation around Stage VIII [74]—a class of genes missed by Hotspot (Fig. S13); and SLOPER module 4 (Figure 5E, bottom right) captures marker genes for early elongated spermatides (ES). See Supplement for more details and biological interpretation of all SLOPER modules considered. A zoom-in analysis of a single tubule (Fig. 5F) further highlights the spatial–temporal coherence of these modules: the mean gradients and resulting module scores for SLOPER modules 6 → 1 → 2 form a clear outward-to-inward progression, mirroring the inward maturation of spermatids from the tubule boundary toward the lumen.

Overall, our results demonstrate that SLOPER’s gradient-based representation not only recovers developmentally ordered gene modules but also captures their spatial colocalization patterns, enabling a unified reconstruction of the spatiotemporal trajectory of spermatogenesis which existing approaches fail to resolve.

## 4 Discussion

We introduce SLOPER, a deep learning algorithm for learning spatial gradients from sparse and spatially diffuse ST data. We derive an inhomogeneous Poisson point process (IPPP) model of spatial gene expression, and we propose a score matching method to learn spatial gradients of expression for individual genes. We derive a novel annealed Langevin dynamics model that leverages the learned spatial gradients to enhance the spatial coherence and specificity of observed gene expression measurements. We demonstrate that the SLOPER-inferred spatial gradients and enhanced gene expression are broadly useful for characterizing tissue organization, spatial gene expression patterns, and axes of continuous variation across different biological systems and ST technologies. We envision that the broad generality of our IPPP and score matching frameworks can be readily extended in several directions including: identification of cell-cell and regulatory interactions mediated by local gradients [75–79]; modeling the spatial dynamics of RNA splicing [80–83]; diffusion-based simulation [84–86] and alignment of multiple ST slices across 3-D or temporal axes [87–90]; and the analysis of intra-cellular spatial gradients [91, 92]. Ultimately, the spatial gradients learned by SLOPER provide a versatile tool for characterizing spatial variation in gene expression from spatial sequencing data.

## Acknowledgements

This work is partially supported by NIH P50 MH094268 (D.G. and L.Y.) and NSF Award 2124230 (A.W., D.G., and L.Y.).

## Notes

### Competing Interest Statement

The authors have declared no competing interest.

