## Supplementary Material for "Mapping spatial gradients in spatial transcriptomics data with score matching"

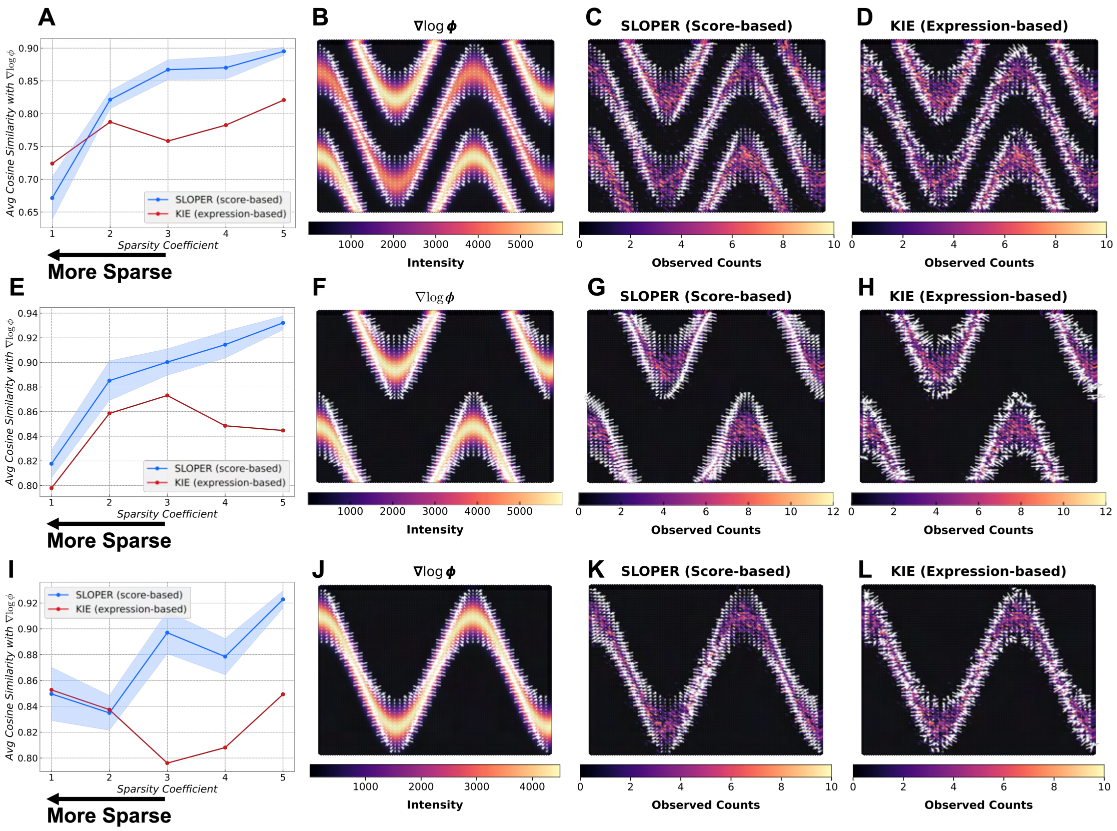


Figure S1: **Benchmark of SLOPER and KIE on Additional IPPP simulation.** For panel A–D, the underlying intensity function was generated using$\{w_l\} = \{0.75,\ 1.0,\ 1.0\}$, $\{b_l\} = \{0,\ 1.5,\ -1.5\}$, $\tau = 0.2$ For panel E–H, the underlying intensity function was generated using $\{w_l\} = \{0.0,\ 1.0,\ 1.0\}$, $\{b_l\} = \{0,\ 1.5,\ -1.5\}$, $\tau = 0.2$. For panel I–L, the underlying intensity function was generated using $\{w_l\} = \{0.75,\ 0.0,\ 0.0\}$, $\{b_l\} = \{0,\ 1.5,\ -1.5\}$, $\tau = 0.2$. For panels B-D, the gradient field is visualized on the subset of spatial locations $\{\mathbf{c}_k\}$ where $\phi(\mathbf{c}_k)$ exceeds the 35% quantile of $\{\phi(\mathbf{c}_k)\}$ to focus on regions with relatively high intensity. For panel F–H and J–L, the gradient field is visualized on the subset of spatial locations $\{\mathbf{c}_k\}$ where $\phi(\mathbf{c}_k)$ exceeds the 18% quantile of $\{\phi(\mathbf{c}_k)\}$ for the same reason. **(A)** Average cosine similarity between the estimated and ground-truth $\nabla \log \phi$ across sparsity coefficients $c$. Shaded regions denote the standard deviation across five random seeds for SLOPER. **(B)** Ground-truth intensity function $\phi$ and the gradient of its log $\nabla \log \phi$. **(C)** SLOPER estimates of $\nabla \log \phi$ (white arrows) overlaid on the simulated Poisson counts. **(D)** Same as (C) but for KIE. **(E)–(H)** Same as (A)–(D) but for the second IPPP hyperparameter configuration. **(I)–(L)** Same as (A)–(D) but for the third IPPP hyperparameter configuration.

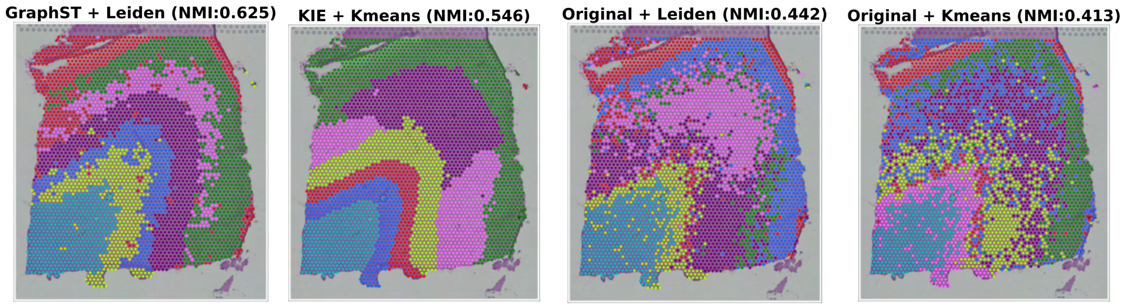


Figure S2: **Domains Identified by GraphST, KIE and original Expression.** Domains are colored according to their closest matching reference annotation the same way as Fig. 3 in the main manuscript.

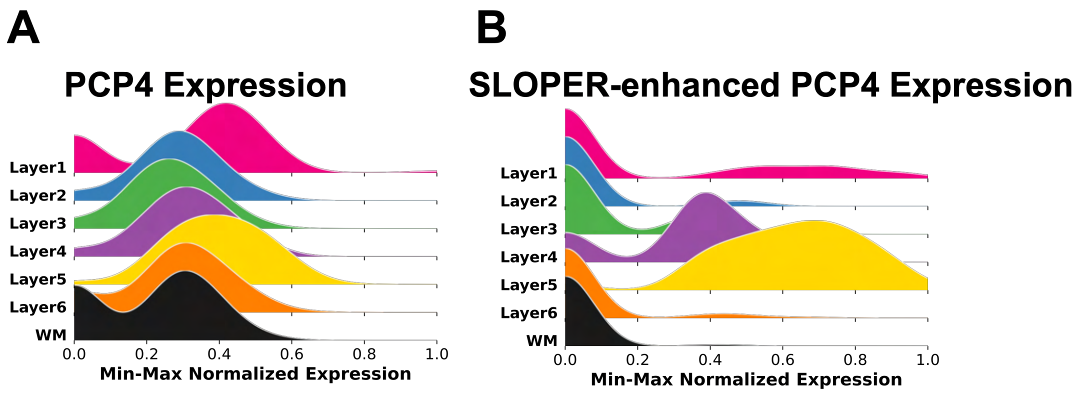


Figure S3: **Normalized *PCP4* before and after Langevin-enhancement. (A)** Left: $[0, 1]$-normalized expression of *PCP4* across cortical layers. **(B)** spatial expression pattern of *PCP4* in the original data.

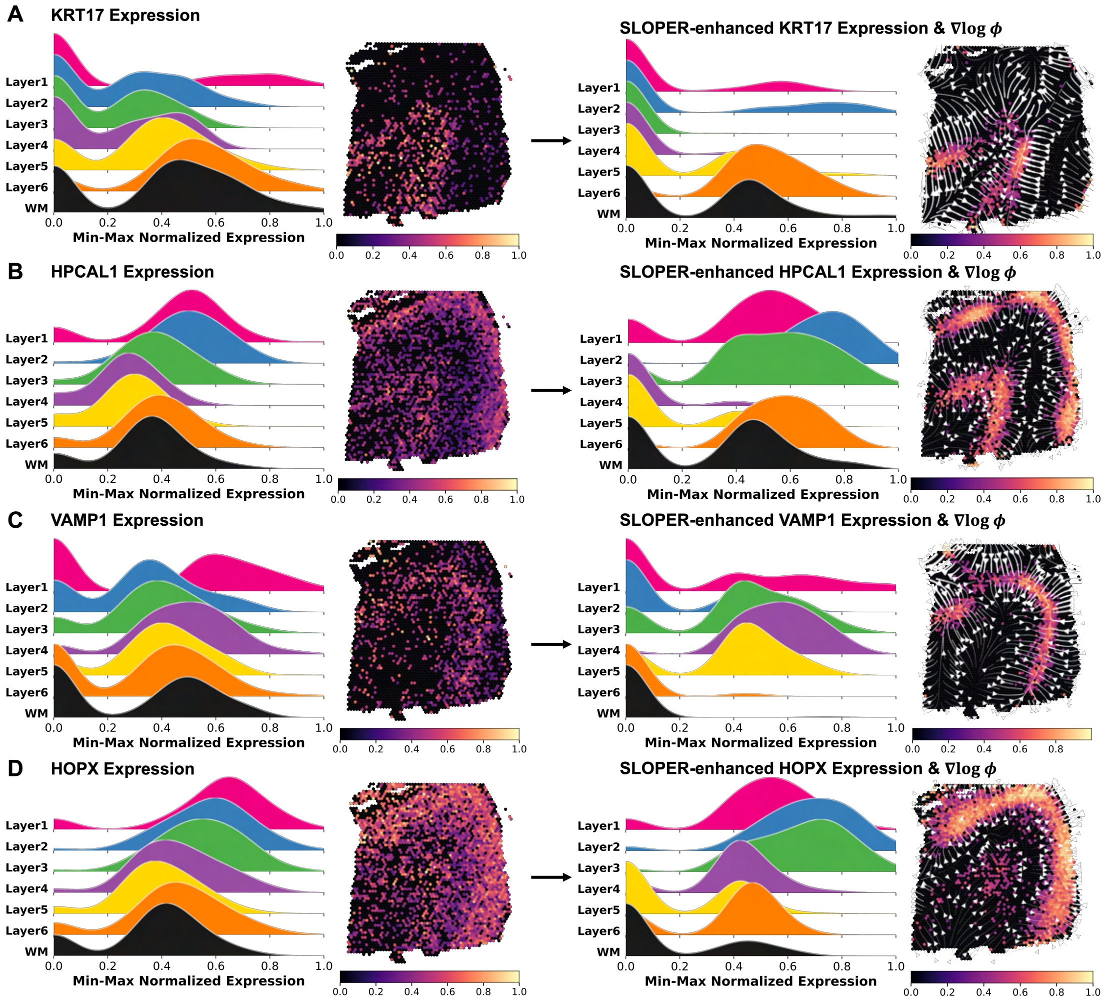


Figure S4: **SLOPER-enhanced expression sharpens layer-specific marker localization across multiple genes. (A)** Left: $[0, 1]$-normalized expression of *KRT17* across cortical layers and its spatial expression pattern in the original data. Right: Same as left panel but for SLOPER-enhanced expression of *KRT17*. **(B)** Same as (A) for *HPCAL1*. **(C)** Same as (A) for *VAMP1*. **(D)** Same as (A) for *HOPX*.

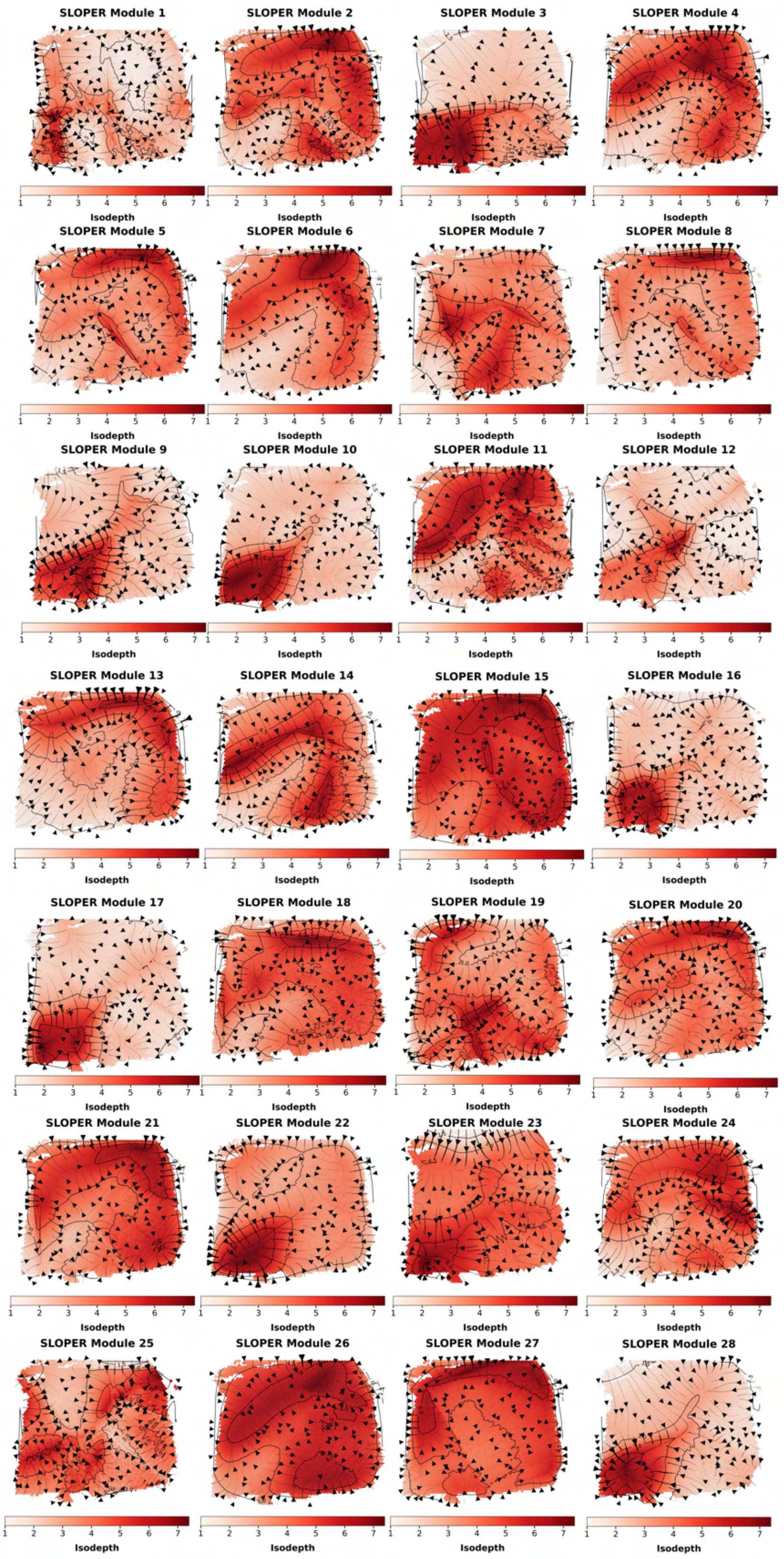


Figure S5: **Gene-module–specific isodepth inference in the DLPFC.** We identified 28 gene modules (Methods) and inferred an isodepth coordinate for each module following Supplementary Section S5. For every module, we computed the corresponding local isodepth gradient with Taylor expansion approximation. The colormap shows the exponential-scaled isodepth, where we normalize the isodepth value to the range [0, 1], and then take the square of its exponential to increase visual contrast.

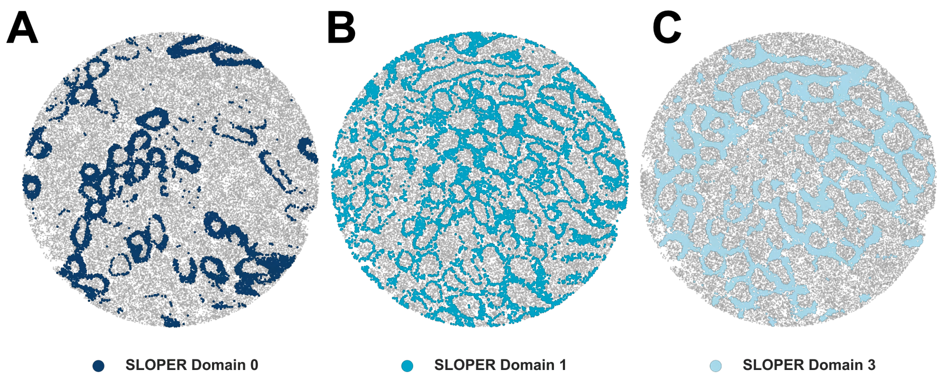


Figure S6: **Visualization of selected SLOPER domains over the Slide-seq spermatogenesis tissue.** Each panel highlights one SLOPER domain (0, 1, or 3) against a gray background to emphasize its spatial organization. **(A)** SLOPER domain 0. **(B)** SLOPER domain 1. **(C)** SLOPER domain 3.

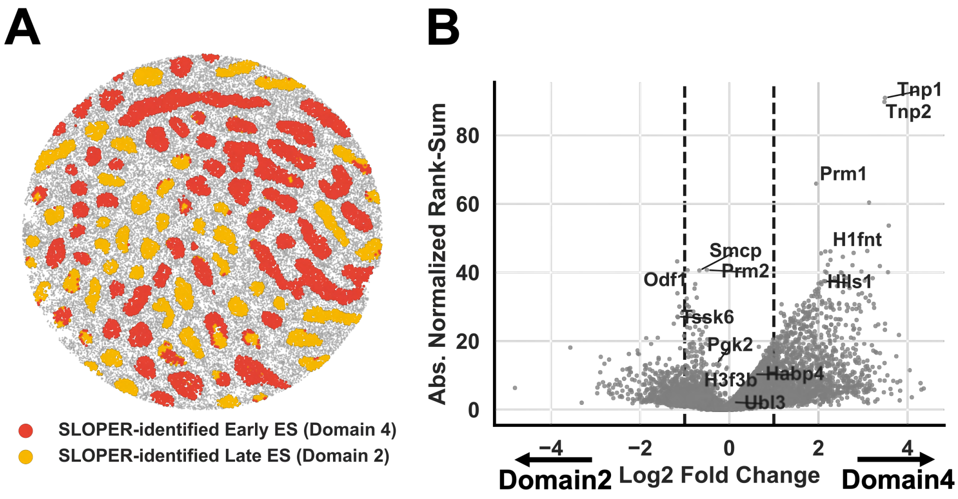


Figure S7: **Differential expression analysis of SLOPER domain 2 and 4. (A)** SLOPER-identified domains 2 and 4 used for differential expression analysis. **(B)** $Log_2$ fold change versus absolute standardized Wilcoxon rank-sum statistic between domains 2 and 4.

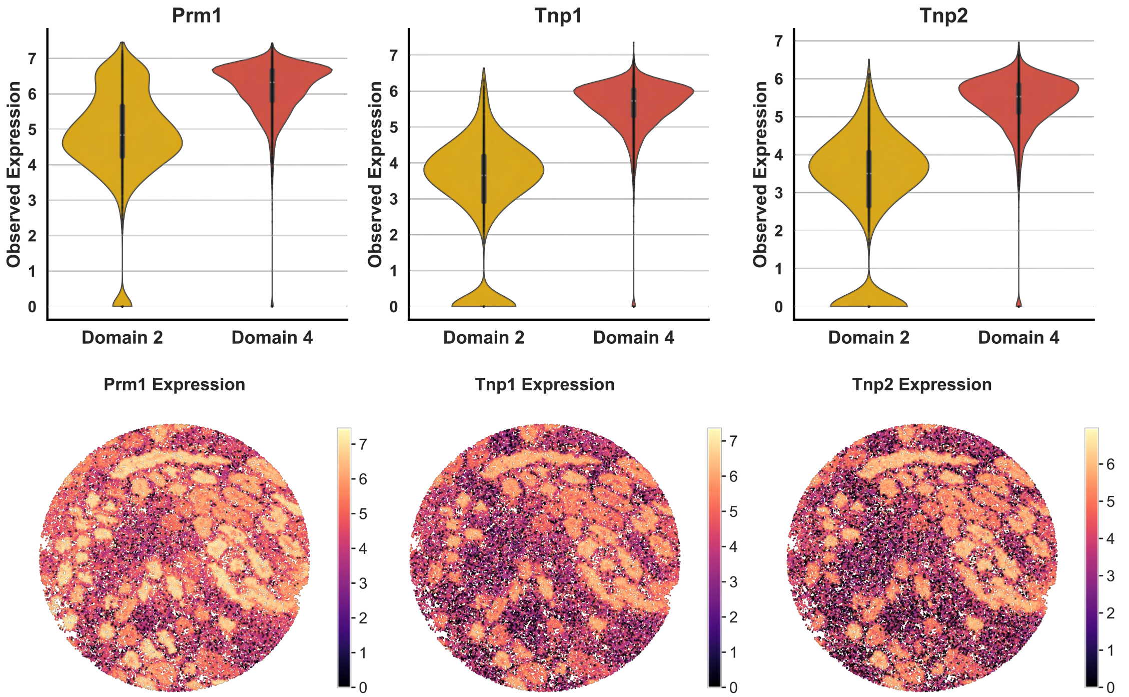


Figure S8: **Expression patterns of marker genes *Prm1*, *Tnp1*, and *Tnp2*.** The top panels show log-normalized expression levels (log-counts per 10,000) across SLOPER domains 2 and 4. The bottom panel highlights their spatial distribution.

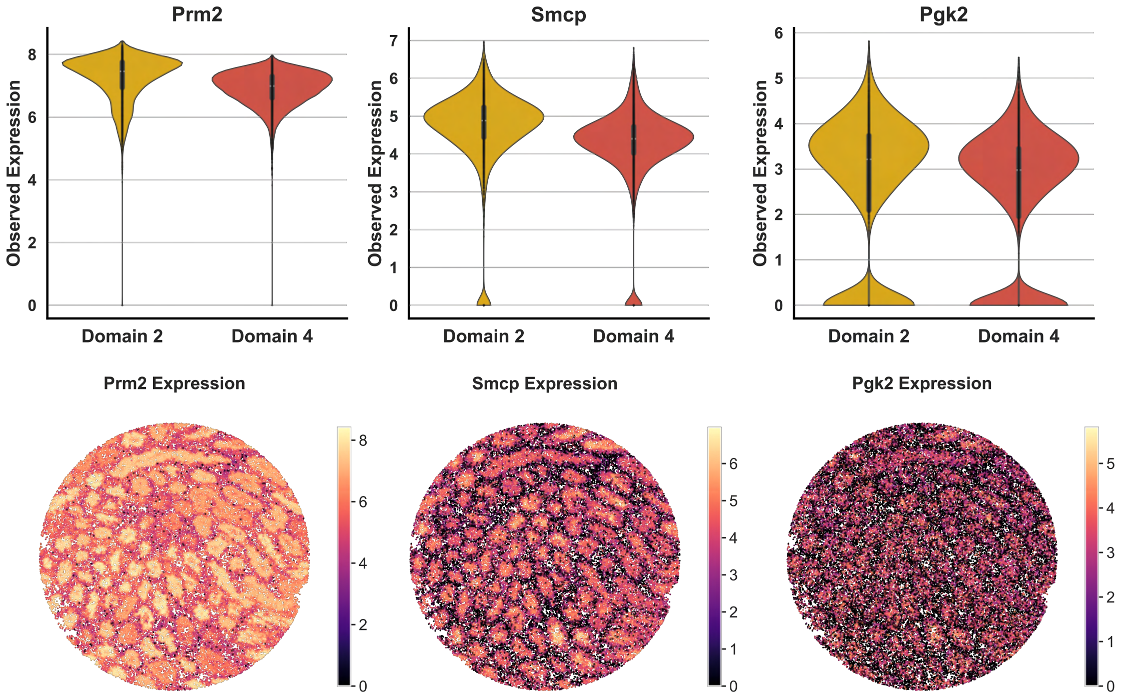


Figure S9: **Expression patterns of marker genes *Prm2*, *Smcp*, and *Pgk2*.** The top panels show log-normalized expression levels (log-counts per 10,000) across SLOPER domains 2 and 4. The bottom panel highlights their spatial distribution.

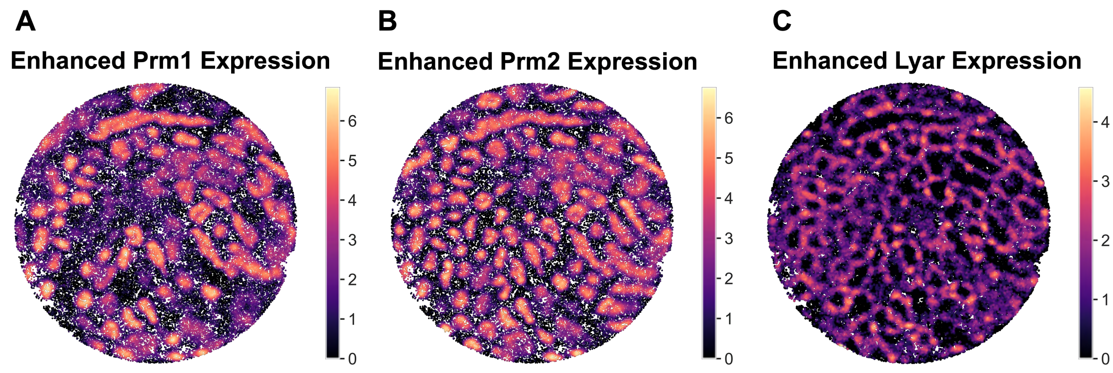


Figure S10: **SLOPER-enhanced spatial expression of *Prm1*, *Prm2*, and *Lyar*.**

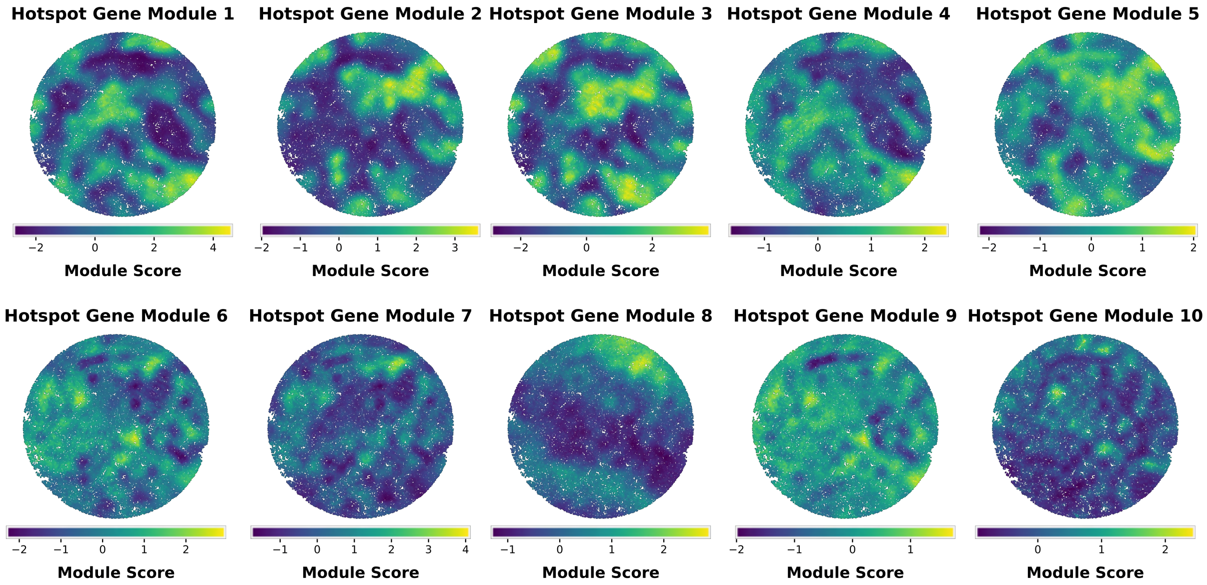


Figure S11: **Hotspot gene modules.** For each module, the colormap represents the per-cell Hotspot module score.

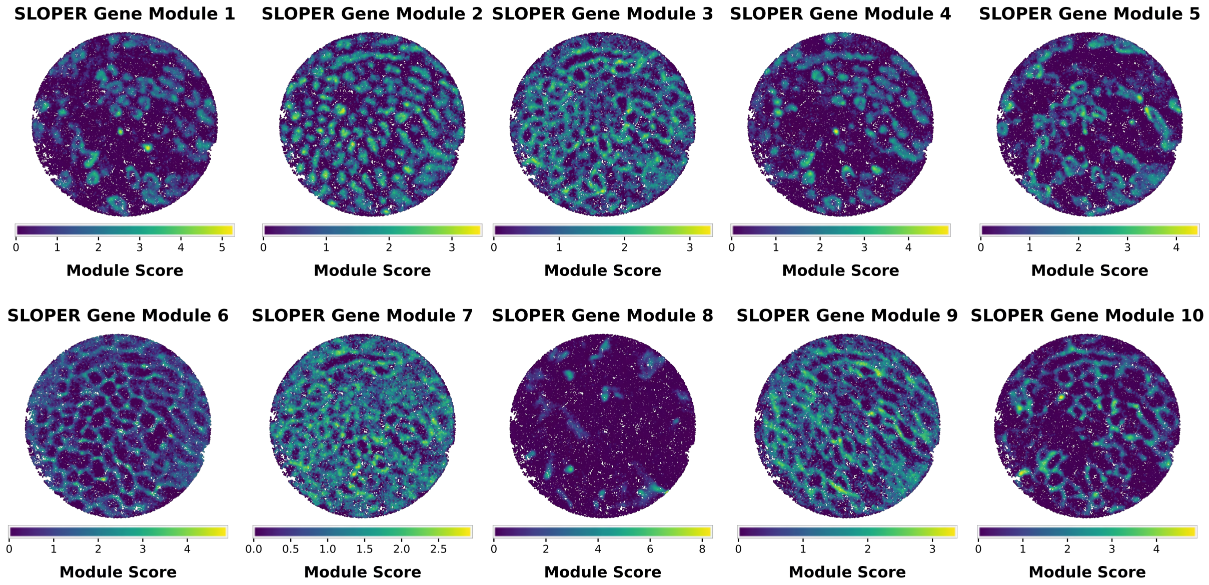


Figure S12: **SLOPER gene modules.** For each module, the colormap represents the per-cell SLOPER module score. All module scores are obtained using annealed Langevin dynamics with step size $\epsilon = 0.03$ and $H = 500$, except for Module 6, for which we use $\epsilon = 0.01$ and $H = 50$ due to the substantially larger gradient norms of the genes in this module.

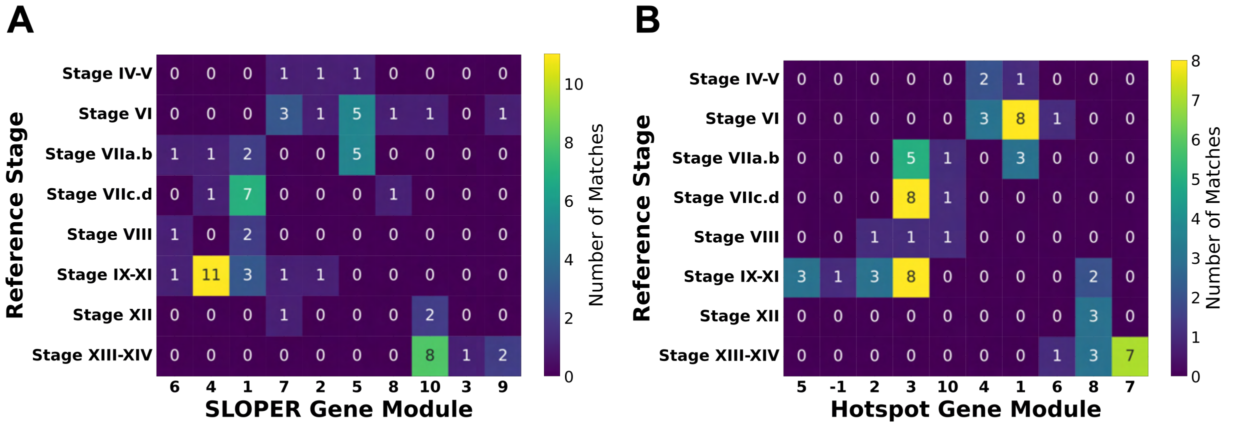


Figure S13: **Mapping of SLOPER and TESTIS modules to spermatogenesis stages.** Each heatmap shows the number of genes that are simultaneously assigned to a spermatogenic stage (x-axis) and a gene module (y-axis). The total number of genes analyzed is 67. **(A)** Mapping based on SLOPER modules. **(B)** Mapping based on Hotspot modules. Among the 500 genes, Hotspot return 10 modules and one additional module (Hotspot Module "$-1$") that contains a small number of unmatched genes based on its algorithm criterion. When taking the intersection of Module "$-1$" genes with the known biological markers, there is one remaining gene– *Prm2*, an important marker for spermatogenesis. This result indicates that Hotspot may miss some biological significant signal in its module identification process. Besides, Hotspot module 9 is missing in this plot since it has no intersection with the known marker genes.

#### SLOPER Modules — Gene Lists

- **Module 1:** *1700012B07Rik, 1700013D24Rik, 1700013G24Rik, 1700015F17Rik, 1700030J22Rik, 1700056E22Rik, 1700074P13Rik, 1700093K21Rik, 1700129C05Rik, 4930578I06Rik, 4931409K22Rik, Akap3, Allc, Aqp7, Arih1, Asns, Bag5, BC049762, Cabyr, Ccdc101, Ccdc183, Ccdc185, Ccdc63, Ccin, Ccser2, Cdkl3, Cdyl, Clip4, Cnbd2, Coro2a, Creb3l4, Cylc2, Cypt12, Cypt4, Desi1, Dgat2, Dpysl3, Erich3, Fam221b, Fhl5, Gm4275, Hils1, Kif2b, Knop1, Lgals8, Lkaaear1, Map7, Micalcl, Nabp1, Pex5l, Prss39, Psmc2, Rnf151, Rnf4, Ropn1, Scp2d1, Sh3d21, Slc16a7, Spag4, Spata21, Sqrdl, Tex29, Tmco2, Tmco5, Tmem144, Tmigd3, Tor1aip1, Trim36, Trim42, Trim80, Tssk1, Ttc23l, Tulp2.*

- **Module 2:** *1110017D15Rik, 1700001O22Rik, 1700003E16Rik, 1700009N14Rik, 1700012A03Rik, 1700016H13Rik, 1700027A15Rik, 1700029H14Rik, 1700031M16Rik, 1700034O15Rik, 4430402I18Rik, 4930571K23Rik, Acsl1, Actg2, Agpat6, Akap1, Akap4, Ankef1, Aqp11, Atp1b3, Bcl2l14, Cabs1, Ccdc67, Ccdc91, Cdrt4, Cep85l, Clpb, D830044I16Rik, Ddi1, Dnajb8, Dnajc5b, Fam217a, Fam71f2, Fxr1, Golph3, H2afb1, Habp4, Hk1, Iqcf3, Klhl10, Lpcat2b, Lrrc57, Luzp1, Mtfr1, Nbr1, Nt5c1b, Osbp2, Oxct2a, Phkg2, Pmfbp1, Prm2, Proca1, Prr30, Pwwp2b, Smcp, Spata18, Spata3, Spem1, St6galnac2, Tesk1, Tppp2, Tssk6, Ttc24, Zbbx.*

- **Module 3:** *1700011E24Rik, 1700028J19Rik, Ccdc173, Ccdc176, Ccdc181, Ccdc38, Cep126, Cep63, Cfap36, Chrac1, Dkkl1, Ggnbp2, Gkap1, Hdgfl1, Larp7, Ldha, Lrrc34, Lrrcc1, Lyar, Mns1, Nasp, Nlrp14, Pebp4, Phospho2, Ppp2r5c, Ppp3r2, Psmc3, Rbakdn, Rnf32, RP23-340E6_2, Rsph6a, Spag17, Spata1, Syce1, Tbpl1, Tcp1, Tdrd6, Tmbim7, Tmem97, Trafd1, Ttc25, Txnrd3, Usp47, Usp8, Vdac2, Wbp11.*

- **Module 4:** *1700016C15Rik, 1700019N19Rik, 1700026L06Rik, 1700034E13Rik, 4921507P07Rik, 4921530L21Rik, 4930407I10Rik, 4930503B20Rik, 4930557A04Rik, Actrt2, Actrt3, Aif1, Asb17, Atp1a4, BC049635, BC100451, Calr3, Capza3, Ccdc54, Cdc34, Cdca2, Chchd3, Chn2, Cst13, Cst8, Cstl1, Dnajb7, Fam187b, Fam71a, Fam71b, Fam71d, Fam71f1, Galntl5, Gm10638, Gpd2, H1fnt, Hmgb4, Hspa1l, Ipo5, Iqcf1, Mgat4e, Otub2, Pcmt1, Pdpk1, Prm1, Prss37, Prss52, Prss55, Rnf133, Samd4, Slfnl1, Spata19, Spata32, Spata6, Spcs2, Sppl2c, Tex37, Tfam, Tnp1, Tnp2, Trim17, Trp53tg5, Tssk2, Tuba8, Txndc2.*

- **Module 5:** *1700001J03Rik, 1700001L19Rik, 1700001P01Rik, 1700007K09Rik, 1700011A15Rik, 1700015E13Rik, 1700016G14Rik, 1700028P14Rik, 4930579F01Rik, Actl7b, Ankrd7, Bbx, BC051628, Best1, Cast, Ccdc116, Ccdc81, Ccdc89, Ccer1, Clip1, Efcab9, Fam170b, Fam209, Ftmt, Gtsf1l, Gykl1, Kcne3, Klf4, Lmnb2, Lyzl1, Lyzl6, Slc22a14, Slc35g3, Sox5, Spaca1, Spaca3, Spaca4, Spag6l, Spata31, Spata45, Tbc1d23, Tekt2, Tex21, Tex33, Tex36, Tmc7, Tsc22d4.*

- **Module 6:** *Agt, Akr1cl, Amhr2, Ccp110, Clu, Cmss1, Cyp17a1, Fabp3, Gm14244, Gstm1, Hexb, Hnrnpa2b1, Hsd3b6, Hspa5, Lars2, Lcn2, Mphosph8, mt-Cytb, mt-Nd1, mt-Nd4, mt-Rnr1, mt-Rnr2, Ncl, Psma8, Sycp1, Sycp2, Sycp3, Tex101, Tex15, Tpr.*

- **Module 7:** *1110032A03Rik, 1700012B09Rik, 1700020D05Rik, 1700086L19Rik, 1700123L14Rik, 4922502D21Rik, Acrbp, Adad1, Adam3, Adam32, Arl4a, Asap1, Asrgl1, BC051142, Ccdc175, Ccdc30, Cep290, Cetn1, Cfap45, Clgn, Cmtm2a, Cmtm2b, Creld2, Dmrtb1, Dnaaf1, Dyrk3, Efcab6, Eif5b, Elof1, Enkur, Ggnbp1, Gk2, Gm11837, Gm128, Gsg2, Gtsf1, Hrasls, Lancl1, Lrrc27, Lypd4, March11, Nphp1, Oscp1, Pabpc1, Papolb, Prdx6b, Pvrl3, Sccpdh, Sucla2, Tekt1, Tekt4, Tex40, Tsnaxip1, Txnrd1, Ube2d2b.*

- **Module 8:** *Acrv1, Aldh1a1, Ccdc105, Cd109, Cfap58, Cyp11a1, Dnah2, Dnajc9, Fsip1, Gm136, Gm6588, Gm8251, Gphn, Hemgn, Lrrc6, Lzts2, Mageb18, Malat1, Prss46, Prss51, Prss58, Ptgds, Saxo1, Star.*

- **Module 9:** *1700003M02Rik, 1700010I14Rik, 1700018B24Rik, AA467197, Camk1d, Catsper2, Ccdc113, Ccdc39, Ccdc96, Cct7, Cenpe, Cenpv, Ddx25, Dydc1, Efcab5, Erich2, Fbp1, Golga4, Gtf2a1l, Hsp90aa1, Hspa2, Larp1b, Mtch2, Nol8, Pabpc2, Prps1l1, Rbbp6, Spag6, Spink2, Stk11, Stk33, Supt20, Tcte1, Zpbp2.*

- **Module 10:** *1700010B08Rik, 2700049A03Rik, 4932431P20Rik, 4933416C03Rik, Abcf1, Ankrd54, Aprt, Atr, Atxn7l3b, Aurka, Calm1, Calm2, Cdc42ep3, Cenpf, Cers3, Cfap97, Cox4i1, Crat, Dcaf7, Diablo, Dnah8, Dnaic2, Fam160b2, Fkbp4, Flywch1, Fsip2, Gm12648, Hsp90b1, Ik, Kif3b, Kpnb1, Ldhal6b, Mllt10, Mum1, Ndufaf3, Nipbl, Nubp2, Pabpc6, Piwil1, Psip1, Rb1cc1, Rpa1, Rsph1, Senp2, Serbp1, Setx, Shcbp1l, Slc2a3, Smarca2, Smc4, Smc6, Spata16, Ssh2, Tchp, Tomm22, Trim37, Tuba3a, Tuba3b, Usp1, Zfp318, Zmynd10.*

### S1 SLOPER neural network architecture and training

For all experiments, we use a fully connected network containing two residual blocks. Each residual block applies two linear layers with an activation in between and adds the output back to the input, followed by a final activation; that is,

$$\text{ResBlock}(x) = a\big(x + W_2\, a(W_1 x)\big),$$

where $a(\cdot)$ denotes the nonlinearity. Each linear layer within the block uses 64 hidden units, and we employ the `softplus` activation function. Networks are trained using the Adam optimizer with learning rate $10^{-3}$.

For the **DLPFC** dataset, we train for 3000 iterations and apply a `StepLR` scheduler with step size 300. We do not use Fourier positional features for this dataset due to the relatively smooth spatial expression patterns.

For the **mouse testis** dataset, we train for 10000 iterations with a `StepLR` step size of 2000. To capture the complex testicular spatial structure, we augment the input coordinates with 64 random Fourier features of the form

$$[\sin(Bx),\ \cos(Bx)], \qquad B \sim \mathcal{N}(0, \sigma^2 I),$$

using $\sigma = 3$.

### S2 Annealed Langevin dynamics

#### S2.1 Hyperparameter selection

The hyperparameters for the annealed Langevin dynamics are chosen according to the following principles.

1. **Step size $\epsilon$.** The step size should balance the magnitude of the learned SLOPER gradient: it must be large enough for the particle to make steady progress across the tissue domain, yet small enough to avoid instability or overshooting.

2. **Number of iterations $H$.** Given a fixed $\epsilon$, the number of iterations must be sufficiently large for particles to reach their attractors and reveal the underlying spatial structure. However, $H$ should not be chosen too large; otherwise, the enhanced feature becomes overly singular (e.g., a very thin and crisp trajectory), which may distort gene-expression enhancement.

In practice, we observe that the average norm of the learned SLOPER gradients in the Testis dataset is substantially larger than that in DLPFC. Consequently, we adopt smaller values of both $\epsilon$ and $H$ for Testis to maintain stable Langevin trajectories.

Following these principles:

- **DLPFC:** For each gene, we set $\epsilon = 0.15$ and $H = 800$. The resulting enhanced features are used for clustering spots.

- **Testis:** For each gene, we set $\epsilon = 0.03$ and $H = 500$. The resulting enhanced features are used for clustering cells.

In addition, for each gene, the number of initial coordinates $N$ is set to be equal to the total raw counts of the gene across the spots. For all analyses, we set annealing rate $\alpha = 0.999$.

#### S2.2 Boundary correction

To ensure that all simulated transcripts remain within the tissue boundary, we implement a boundary-constrained Langevin sampling procedure based on a signed distance field (SDF) representation of the tissue polygon.

**Signed distance field rasterization.** We first rasterize the tissue boundary, represented as a polygon $\Omega \subset \mathbb{R}^2$, into a high-resolution grid to obtain a signed distance field $\psi(x)$ that records, for each spatial location, its Euclidean distance to the boundary (negative inside, positive outside). The rasterization uses exact point-in-polygon queries on a regular lattice, followed by Euclidean distance transforms to compute the interior and exterior distances in world units. The resulting SDF provides both a continuous approximation of the tissue mask and a differentiable proxy for boundary gradients.

**GPU SDF projector.** The precomputed SDF is transferred to the GPU and wrapped in an `SDFProjector` module that supports three key operations: (i) bilinear interpolation of $\psi(x)$ to obtain approximate signed distances at arbitrary coordinates, (ii) finite-difference estimation of spatial gradients $\nabla\psi(x)$ in world units, and (iii) a Newton-like projection step that maps any point lying outside the domain ($\psi(x) > 0$) back to the boundary ($\psi(x) \approx 0$) and then a small step inward along $-\nabla\psi(x)$ to ensure interior placement. This projector provides efficient, differentiable access to boundary geometry entirely on the GPU.

**Projected Langevin dynamics.** We integrate the learned score field $f_{\theta^*}(x)$ with the SDF projector to perform Langevin sampling constrained to the tissue domain. At each iteration,

$$\mathbf{s}^{(t+1)} = \mathbf{s}^{(t)} + \frac{\varepsilon^2}{2}\, f_{\theta^*}(\mathbf{s}^{(t)}) + \varepsilon\, \sigma_t\, z^{(t)}, \qquad z^{(t)} \sim \mathcal{N}(0,1), \quad \sigma_t = \alpha^t,\ \alpha < 1, \tag{S1}$$

and every few steps the updated coordinates are projected using the SDF-based Newton correction to ensure $x^{(t)} \in \Omega$. This "projected Langevin" scheme combines the flexibility of stochastic diffusion sampling with strict geometric constraints imposed by the tissue boundary, enabling realistic simulation of transcript distributions even for highly non-convex tissue shapes.

#### S3 Simulation set-up

To assess the accuracy of score matching in recovering the gradient of the log intensity, we benchmarked **SLOPER** against the **kernel intensity estimator (KIE)** [63] on simulated IPPP data with known ground truth. We considered an IPPP with intensity function $\phi(s)$, assuming $u(s) = 1$ for all $s \in T$. The spatial domain was defined as $T = [-1.5, 1.5] \times [-1.5, 1.5]$, and the true intensity function was given by:

$$\phi(x, y) = 1000c \cdot \sum_{l=1}^{3} w_l \exp\left(-\frac{(y - \mu_l(x))^2}{2\tau^2}\right)$$

$$\nabla \log \phi(x, y) = \frac{1000c}{\tau^2 \phi(x, y)} \begin{bmatrix} \sum_{l=1}^{3} w_l (y - \mu_l(x)) \mu_l'(x) \exp\left(-\frac{(y-\mu_l(x))^2}{2\tau^2}\right) \\ -\sum_{l=1}^{3} w_k (y - \mu_l(x)) \exp\left(-\frac{(y-\mu_l(x))^2}{2\tau^2}\right) \end{bmatrix}$$

where each curve $\mu_l(x) = \sin(\pi x) + b_l$ defines a wavy Gaussian ridge, $\mu_l'(x) = \pi \cos(\pi x)$, $w_l$ and $b_l$ are the weight and offset parameters for the Gaussian ridge, and $c$ is a sparsity coefficient controlling overall density (smaller $c$ yields sparser data).

We evaluated both estimators over multiple parameter settings $\{w_l, b_l\}_{l=1:3}$ and sparsity levels $c \in \{1, 2, 3, 4, 5\}$. To simulate the data, the domain $T$ was divided uniformly into 10,000 spots $\{C_k\}$. For each spot, counts $a_k \sim \text{Poisson}(\tilde{\Lambda}(C_k))$ were sampled, with $\tilde{\Lambda}(C_k) = \frac{9}{10000}\phi(\mathbf{c}_k) \approx \Lambda(C_k)$, where $bc_k$ denotes the centroid of cell $C_k$.

#### S4 Kernel Intensity Estimator

Let $k : \mathbb{R}^2 \to \mathbb{R}_+$ be a spherically symmetric kernel that integrates to 1 and $h$ be the bandwidth. Let the dimension-scaled kernel be $k_h(z) = h^{-2}k(\frac{z}{h})$. Given the transcript locations $\{x_i\}_{i=1:N}$. The Jone-Diggle kernel estimator at $x \in \Omega$ [63] is

$$\hat{\lambda}_h(x) = \sum_{i=1}^{N} \frac{k_h(x - x_i)}{w_h(x_i)}$$

where $w_h(x) = \int_\Omega k_h(u - x)du$.

Jone-Diggle kernel estimator is **mass preserving**:

$$\int_\Omega \hat{\lambda}_h(x)dx = N$$

Likelihood cross-validation is used for optimal bandwidth selection. Define

$$LCV(h) = \sum_{i=1}^{N} \log \hat{\lambda}_h^{(-i)}(x_i) - \int_\Omega \hat{\lambda}_h(x)dx$$

where $\hat{\lambda}_h^{(-i)}(x_i) = \sum_{j \neq i} \frac{k_h(u - x_i)}{w_h(x_i)}$. Then $h^* = \arg\min_h LCV(h)$.

The kernel intensity estimator can also be adapted to the situation when only the spot by counts matrix is available instead of the transcript locations. We simply consider the multiplicity at each spot where the multiplicity equals the count at the spot. Given bandwidth $h$ and spot locations-counts pairs $\{(s_i, c_i)\}_{i=1:n}$, we now have

$$\hat{\lambda}_h(x) = \sum_{i=1}^{n} \frac{c_i \cdot k_h(x - s_i)}{w_h(s_i)}$$

Note that the mass-preserving property is maintained.

For the optimal bandwidth selection, we can consider the same cross-validation framework with a leave-one-count-out scheme where

$$\hat{\lambda}_h^{(-i)}(s_i) = \frac{1}{w_h(s_i)} \sum_{j \neq i} (c_j k_h(s_i - s_j) + (c_i - 1)k_h(0))$$

**Learning Variation of the True Expression: Kernel Intensity Estimator** We assume the Poisson measurement model

$$\lambda(x) \;=\; u(x)\,\phi(x), \qquad x \in \Omega,$$

where $u(x) > 0$ is the local UMI "effort" (library size / sampling rate) and $\phi(x)$ is the true expression we want to estimate.

Let $k_h(\cdot)$ be a symmetric kernel with bandwidth $h > 0$ (e.g. $k_h(z) = h^{-2}k(z/h)$ in 2D), and let $\{x_i\}_{i=1}^N$ be observed transcript locations (a point pattern). Define the *UMI-weighted boundary factor*

$$w_h^{(u)}(x) \;=\; \int_\Omega u(v)\,k_h(v-x)\,dv.$$

**Point-pattern version.** Define the *UMI-aware Jones–Diggle estimator* for $\phi$:

$$\hat\phi_h(x) \;=\; \sum_{i=1}^N \frac{k_h(x-x_i)}{w_h^{(u)}(x_i)}.$$

*Mass preservation:*

$$\int_\Omega \hat\lambda_h(x)\,dx = \int_\Omega u(x)\hat\phi_h(x)\,dx = \sum_{i=1}^N \frac{1}{w_h^{(u)}(x_i)} \int_\Omega u(x)k_h(x-x_i)\,dx = \sum_{i=1}^N 1 = N.$$

*Likelihood cross-validation (LCV):*

$$LCV(h) \;=\; \sum_{i=1}^N \log\big(\hat\lambda_h^{(-i)}(x_i)\big) \;-\; \int_\Omega \hat\lambda_h(x)\,dx,$$

where

$$\hat\lambda_h^{(-i)}(x) = u(x)\,\hat\phi_h^{(-i)}(x), \qquad \hat\phi_h^{(-i)}(x_i) \;=\; \sum_{j\neq i} \frac{k_h(x_i-x_j)}{w_h^{(u)}(x_j)}.$$

The optimal bandwidth is chosen as

$$h^\star \;=\; \arg\max_h \; LCV(h).$$

**Spot–count version.** Suppose we have spot locations $\{s_j\}_{j=1}^n$ with counts $\{c_j\}_{j=1}^n$. Treating each spot as $c_j$ co-located points gives

$$\hat\phi_h(x) \;=\; \sum_{j=1}^n \frac{c_j\,k_h(x-s_j)}{w_h^{(u)}(s_j)}.$$

In practice, we approximate $\mathrm{w}_h^{(u)}(x)$ using a finite sum over observed spot locations:

$$w_h^{(u)}(x) \;\approx\; \sum_{j=1}^n u(s_j)\,k_h(s_j-x)\,\Delta A,$$

where $\Delta A$ denotes the area associated with spot $j$.

***Mass preservation:***

$$\int_\Omega u(x)\hat\phi_h(x)\,dx = \sum_{j=1}^n \frac{c_j}{w_h^{(u)}(s_j)} \int_\Omega u(x)k_h(x-s_j)\,dx = \sum_{j=1}^n c_j \;=\; C,$$

where $C = \sum_j c_j$ is the total count for the gene.

***Leave-one-count-out log pseudo-likelihood:*** At spot $i$, removing one of its $c_i$ co-located counts yields

$$\hat\phi_h^{(-i)}(s_i) \;=\; \sum_{j\neq i} \frac{c_j\,k_h(s_i-s_j)}{w_h^{(u)}(s_j)} \;+\; \frac{(c_i-1)\,k_h(0)}{w_h^{(u)}(s_i)}.$$

Hence

$$LCV(h) \;=\; \sum_{i=1}^n c_i \log\Big(u(s_i)\hat\phi_h^{(-i)}(s_i)\Big) \;-\; \int_\Omega \hat\lambda_h(x)\,dx, \qquad h^\star = \arg\max_h LCV(h).$$

#### S5 Learning isodepth and topographic maps

Suppose $\nabla \log \phi_g = c_g \nabla d$ for some scalar field $d : \mathbb{R}^2 \to \mathbb{R}$ for a subset of genes $g \in \mathcal{M} \subseteq \{1, \ldots, G\}$ and for all spatial locations $(x, y)$ in a domain. That is, the normalized expression function $\log \phi_g$ of each gene $g \in \mathcal{M}$ is a linear function of a scalar field $d(x, y)$ which we call isodepth. This matches the assumption from GASTON [34, 35].

Define $P_{(x,y)} = \frac{1}{|\mathcal{M}|} \sum_{g \in \mathcal{M}} (\nabla \log \phi_g(x, y))(\nabla \log \phi_g(x, y))^T$, or the gradient outer product averaged over all genes $g$. Plugging in $\nabla \log \phi_g = c_g \nabla d(x, y)$ yields:

$$P_{(x,y)} = \frac{1}{|\mathcal{M}|} \sum_{g \in \mathcal{M}} (\nabla \log \phi_g(x, y))(\nabla \log \phi_g(x, y))^T = \left( \frac{1}{|\mathcal{M}|} \sum_{g \in \mathcal{M}} c_g^2 \right) (\nabla d(x, y))(\nabla d(x, y))^T = \widetilde{c^2} (\nabla d(x, y))(\nabla d(x, y))^T, \tag{S2}$$

where we define $\widetilde{c^2} = \left( \frac{1}{|\mathcal{M}|} \sum_{g \in \mathcal{M}} c_g^2 \right)$.

Note that the square root $P_{(x,y)}^{1/2}$ of the gradient outer product matrix is given by

$$P_{(x,y)}^{1/2} = \frac{\sqrt{\widetilde{c^2}}}{||\nabla d(x, y)||} (\nabla d(x, y))(\nabla d(x, y))^T. \tag{S3}$$

The norm $||P_{(x,y)}^{1/2} \mathbf{v}||$ of the square-root $P_{(x,y)}^{1/2}$ of the gradient outer product multiplied by a vector $\mathbf{v} \in \mathbb{R}^2$ is proportional to the (absolute value of the) *directional derivative* of the isodepth $d$ in the direction $\mathbf{v}$:

$$||P_{(x,y)}^{1/2} \mathbf{v}|| = \widetilde{c^2} \, |(v^T)(\nabla d(x, y))| \iff |(v^T)(\nabla d(x, y))| = \frac{1}{\widetilde{c^2}} ||P_{(x,y)}^{1/2} \mathbf{v}|| \tag{S4}$$

**Gene module specific isodepth inference.** For gene-module–specific isodepth inference, we determine the sign sgn $(v^T)(\nabla d(x, y))$ of the directional derivative $(v^T)(\nabla d(x, y))$ at each location $(x, y)$ by computing which of $P_{(x,y)}^{1/2} \mathbf{v}$ and $-P_{(x,y)}^{1/2} \mathbf{v}$ has a higher cosine similarity with the average estimated spatial gradient $\frac{1}{|\mathcal{M}|} \sum_{g \in \mathcal{M}} \nabla \log \phi_g$ over all genes $g \in \mathcal{M}$. This produces a signed directional derivative consistent with the dominant orientation of the gradients within the module.

**Global isodepth inference.** In contrast, when inferring the global isodepth using all marker genes considered (as done in the main text), we do *not* resolve this sign ambiguity. The goal in the global analysis is to identify a coherent *axis* of spatial variation rather than a direction with a prescribed orientation. Because this axis is directionless, choosing a particular sign is unnecessary; we simply adopt one of the equivalent orientations.

Euler's method gives a simple approach for estimating a function $d(x, y)$ given a single value $d(x_0, y_0)$ at a fixed point $(x_0, y_0)$ and a black-box approach to compute the directional derivative of $(\nabla d)^T \mathbf{v}$ in any direction $\mathbf{v}$. Without loss of generality, assume $d(0, 0) = 0$. The first-order Taylor expansion for $d$ at a point $(x, y)$ close to the origin $(0, 0)$ is given by:

$$d(x, y) \approx (\nabla d(0, 0))^T \mathbf{v}, \tag{S5}$$

where $\mathbf{v} = \begin{pmatrix} x \\ y \end{pmatrix}$.

For locations $(x, y)$ that are far from the origin, the Taylor expansion (S5) may not be accurate. We instead create $Q$ equally spaced points $u_0 = (0, 0), u_1 = (\frac{1}{Q} x, \frac{1}{Q} y), u_2 = (\frac{2}{Q} x, \frac{2}{Q} y), \cdots, u_Q = (x, y)$ between the origin $(0, 0)$ and $(x, y)$ and estimate $d(x, y)$ as

$$d(x, y) \approx \sum_{q=1}^{Q} \frac{1}{Q} \big( (\nabla d(u_{q-1}))^T \mathbf{v} \big). \tag{S6}$$

After learning the isodepth function $d(x, y)$, we estimate its spatial gradient $\nabla d(\mathbf{c}_i)$ at each spot $\mathbf{c}_i$ using a local first–order Taylor approximation. For a given spot $\mathbf{c}_i$, let $\{\mathbf{c}_j\}_{j \in \mathcal{N}(i)}$ denote its $k$ nearest neighbors. In a neighborhood of $\mathbf{c}_i$, we approximate $d$ by its first–order Taylor expansion

$$d(\mathbf{c}_j) \;\approx\; d(\mathbf{c}_i) + \nabla d(\mathbf{c}_i)^\top (\mathbf{c}_j - \mathbf{c}_i), \qquad j \in \mathcal{N}(i).$$

Rearranging terms gives

$$d(\mathbf{c}_j) - d(\mathbf{c}_i) \;\approx\; (\mathbf{c}_j - \mathbf{c}_i)^\top \nabla d(\mathbf{c}_i).$$

Define the vectors

$$z_i \;=\; \big(d(\mathbf{c}_j) - d(\mathbf{c}_i)\big)_{j \in \mathcal{N}(i)} \in \mathbb{R}^k, \qquad B_i \;=\; \big(\mathbf{c}_j - \mathbf{c}_i\big)_{j \in \mathcal{N}(i)} \in \mathbb{R}^{k \times 2}.$$

Then the Taylor relations can be written compactly as

$$z_i \;\approx\; B_i \,\nabla d(\mathbf{c}_i).$$

We estimate $\nabla d(\mathbf{c}_i)$ by solving a (weighted) least–squares problem:

$$\widehat{\nabla d}(\mathbf{c}_i) = \arg\min_{\mathbf{v} \in \mathbb{R}^2} \sum_{j \in \mathcal{N}(i)} w_{ij} \left(d(\mathbf{c}_j) - d(\mathbf{c}_i) - (\mathbf{c}_j - \mathbf{c}_i)^\top \mathbf{v}\right)^2,$$

where $w_{ij}$ is a spatial weight that decreases with the distance $\|\mathbf{c}_j - \mathbf{c}_i\|$ (e.g. a Gaussian kernel). This minimization has the closed-form solution

$$\widehat{\nabla d}(\mathbf{c}_i) = (B_i^\top W_i B_i)^{-1} B_i^\top W_i z_i, \quad W_i = \mathrm{diag}(w_{ij}).$$

### S6 DLPFC

We applied SLOPER and KIE to 500 marker genes selected by geneCover [65] without using layer annotations. From the learned SLOPER score, we refined the gene expression features using annealed Langevin diffusion and subsequently performed $k$-means clustering on the enhanced features. The number of clusters was set to $k = 7$ to match the ground-truth domain annotations. For KIE, we applied the same clustering pipeline to its estimated intensity field $\tilde{\phi}_{\text{KIE}}$. We note that we applied standard normalization (per-cell UMI normalization and log-transformation as implemented in Scanpy [61]) to SLOPER-enhanced features, KIE intensities, and the original raw counts before clustering. As additional baselines, we also applied both the $k$-means and Leiden algorithms directly to the processed original gene expression matrix using the same 500 geneCover marker genes. For comparison with a state-of-the-art spatial domain detection method, we evaluated GraphST [15] following the method's' default pipeline, where domains were obtained by applying the Leiden algorithm to the learned embeddings from the top 3000 highly variable genes.

To illustrate this, we extracted the top 15 differentially expressed (DE) genes from each ground-truth layer using the Wilcoxon rank-sum test applied to the original expression matrix (See below). For each DE gene, we quantified how well its expression (both original and Langevin-enhanced) distinguishes the layer from all others by computing the area under the receiver operating characteristic curve (AUROC). For each layer, we then report the mean AUROC across its 15 DE genes. As shown in Figure 3C, the SLOPER-enhanced expression consistently achieves higher average AUROC scores than the original expression across Layers 1–6, indicating that SLOPER increases the discriminative power of gene expression profiles with respect to cortical layer identity.

**Wilcoxon rank-sum test:** For each layer $k$, we identify differentially expressed genes using the Wilcoxon rank–sum test implemented in `scanpy.tl.rank_genes_groups()`. For a given gene $g$, let $A_k$ denote the set of cells in layer $k$ and $B_k$ the set of all remaining cells. Let $a_{i,g}$ denote the expression of gene $g$ in cell $i$, and let $R_{i,g}$ be the rank of $a_{i,g}$ among all cells in $A_k \cup B_k$. The Wilcoxon rank–sum statistic for gene $g$ in layer $k$ is

$$W_{kg} \;=\; \sum_{i \in A_k} R_{i,g},$$

the sum of ranks within the target layer. Scanpy standardizes this statistic using its null mean and variance (with tie correction), yielding the z–score

$$Z_{kg} \;=\; \frac{W_{kg} - \mathbb{E}[W_{kg}]}{\sqrt{\mathrm{Var}(W_{kg})}},$$

which we refer to as the *DE score*. Large positive values of $Z_{kg}$ indicate that gene $g$ tends to have higher expression in layer $k$ relative to all other layers. For each layer $k$, we perform a one–vs–rest comparison and select the top 15 genes with the highest DE scores.

**AUROC:** To quantify how well each gene distinguishes a given layer from all others, we computed one-vs-rest area under the receiver operating characteristic curve (AUROC) using both the original expression and the Langevin-enhanced values. For each layer $k$, we define a binary indicator

$$y_i^{(k)} = \mathbb{I}\{\ell_i = k\},$$

and treat the expression values $\{a_{i,g}\}_i$ as a continuous feature for distinguishing cells in layer $k$ from all other layers. The AUROC for gene $g$ with respect to layer $k$ is then the area under the ROC curve obtained by thresholding the feature $a_{i,g}$ at all possible values. Equivalently, the AUROC is the probability that a randomly chosen cell from layer $k$ has a higher expression of gene $g$ than a randomly chosen cell from any other layer, with ties counted as one-half:

$$\mathrm{AUROC}_{gk} = \Pr\big(a_{i,g} > a_{j,g} \,\big|\, \ell_i = k,\ \ell_j \neq k\big) + \frac{1}{2}\Pr\big(a_{i,g} = a_{j,g} \,\big|\, \ell_i = k,\ \ell_j \neq k\big).$$

This provides a threshold-free measure of how well the expression of gene $g$ separates layer $k$ from all others. The resulting one-vs-rest AUROC for gene $g$ and layer $k$ is denoted $\mathrm{AUROC}_{gk}$. For each gene, we record $\mathrm{AUROC}_{gk}$ for all layers $k$ and summarize its discriminative power by the *best-layer AUROC*

$$\max_k \mathrm{AUROC}_{gk},$$

along with the corresponding layer achieving this maximum.

#### S7 Mouse testis

**Domains.** Using 500 label-free geneCover markers, we applied both SLOPER and KIE to obtain spatial features and performed $k$-means clustering, comparing results to IRIS [13], GraphST [15], and GASTON [34] ran with their default pipeline. SLOPER delineates all major testicular cell types and reveals potential developmental stage differences among elongated spermatids. Despite the lack of ground-truth annotations, the spatial colocalization of testicular cells provides clear structural cues for identifying distinct cell types [13, 69]. As shown in Fig. 5A, SLOPER accurately resolves the testicular architecture: domain 0 (Fig. S6A) corresponds to round spermatids; domain 1 (Fig. S6B) represents spermatocytes, a granular population colocalizing with spermatids but missed by GraphST, KIE, and GASTON; domains 2 and 4 (Fig. S7A) correspond to distinct stages of elongated spermatids, which IRIS merges into a single domain; and domain 3 (Fig. S6C) captures the interstitial region containing Sertoli and endothelial cells. The ability to recover such morphologically complex structures demonstrates that the gradient-derived features of SLOPER effectively capture fine-scale spatial organization.

**Gene modules.** The overlap between SLOPER gene modules and known spermatogenesis developmental stage markers (Figure S13A) reveals the spatial organization of cells along the spermatogenic cycle. For example, SLOPER modules 3, 9, and 10 have substantial overlap with marker genes for spermatogenesis stages XIII-XIV; e.g., SLOPER module 10 contains *Aurka*, a gene essential for spermatocyte maintenance [72] and meiosis progression [93]. In spermatogenesis stages XIII-XIV, spermatocytes undergo meiotic divisions and produce haploid, round spermatids that will re-enter the spermatogenesis cycle at stage I [71, 73]. Visually, we observe that the corresponding SLOPER modules (e.g. SLOPER module 3, Fig. 5E, top right) have large module scores in thin spatial regions that surround the repeating, tubular structures of the testis (SLOPER domains 2 and 4, Figure S7A). The spatial pattern of these SLOPER module scores suggests that new haploid, round spermatids are produced on the boundary of the tubules where spermatogenesis takes place, which is consistent with previous biological studies on the spatial organization of the testis [94]. Furthermore, SLOPER module 5 has a module score pattern (Figure 5E, top right) that visually aligns with the location of round spermatid cells reported in the original study [69] and also contains some known round spermatid marker genes (e.g. *Actl7b, Gtsf1l*) [95, 96]—consistent with the observation that round spermatids exhibit elevated marker gene expression during Stage VI [71]. Meanwhile, SLOPER module 1 has higher overlap with the subsequent stage VIIc.d, the stage in which round spermatids start to elongate and mature Endo et al. [74], demonstrating that these two SLOPER modules capture a temporal progression. The reduced spatial extent of the SLOPER module 1 score relative to SLOPER module 5 score (Figure 5E, top right and bottom right) in certain round spermatid domains may reflect the biological decrease in round spermatids as they enter elongation [94]. Lastly, SLOPER module 4 has high overlap with stage IX–XI marker genes (e.g. *Prm1*, *Tnp1*, *Tnp2*) and has large module score in SLOPER domain 2 (Figure 5E, bottom left; Figure S7A), indicating that this module represents marker genes for early stage elongated spermatids [94].
